# To See, Not to See, or to See Poorly: Perceptual Quality and Guess Rate as a Function of Electroencephalography (EEG) Brain Activity in an Orientation Perception Task

**DOI:** 10.1101/2020.12.21.423809

**Authors:** Sarah S. Sheldon, Kyle E. Mathewson

**Author notes:** Correspondence concerning this article should be addressed to Sarah S. Sheldon, Department of Psychology, University of Alberta, Edmonton, Alberta, T6G 2R3, Canada.

## Abstract

Detection of visual stimuli fluctuates over time, and these fluctuations have been shown to correlate with time-domain evoked activity and frequency-domain periodic activity. However, it is unclear if these fluctuations are related to a change in guess rate, perceptual quality, or both. Here we determined whether the quality of perception randomly varies across trials or is fixed so that the variability is the same. Then we estimated how perceptual quality and guess rate on an orientation perception task relates to EEG activity. Response errors were fitted to variable precision models and the standard mixture model to determine whether perceptual quality is from a varying or fixed distribution. Overall, the best fit was the standard mixture model that assumes response variability can be defined by a fixed distribution.

The power and phase of 2-7 Hz post-target activities were found to vary along with task performance in that more accurate trials had greater power, and the preferred phase differed significantly between accurate and guess trials. Guess rate and σ were significantly lower on trials with high 2-3 Hz power than low and the difference started around 250 ms post-target.

These effects coincide with changes in the P3 ERP: there was a more positive deflection in the accurate trials vs guesses. These results suggest that the spread of errors (perceptual quality) can be characterized by a fixed range of values. Where the errors fall within that range is modulated by the post-target power in the lower frequency bands and their analogous ERPs.

---

Variations in neural activity give rise to observed variations in our visual perception. (Chaumon & Busch, 2014; Mathewson et al., 2011; Samaha et al., 2020). While there has been a plethora of research into the brain activity that drives this process, there remains noticeable gaps in our understanding of how these processes work. One of the reasons for this might be because investigators have left basic questions about the underlying mechanisms unanswered.

Specifically, and the question the current research will address, does a perceptual representation always form with the same precision or does the quality depend on the state of the neural activity?

Traditionally, visual perception has been studied with two-alternative forced-choice (2- AFC) tasks or similar discrete response paradigms. In these types of paradigms, participants are required to select one out of two or more possible responses. Sometimes they are asked to choose the correct stimulus out of an array of different stimuli or to simply report whether they detected a visual stimulus. While these paradigms are powerful and easy to use, they might not be the best choice for investigating certain aspects of visual perception that are easier to measure with a continuous scale. For example, the question of whether the quality of visual perception varies from trial to trial or has a precision that remains constant for a given level of visibility would be difficult to answer without a way to directly measure the variability of a response, something that cannot be done with categorical data (in regards to the traditional concept of variability; see Kader and Perry (2007) for a discussion on variability in categorical data). To answer this question about the nature of perceptual processes, we chose a task that can measure visual perception on a continuous scale and a model that can quantify perceptual responses in a way that will inform our question (it should be noted that this can be done by using categorical responses (see Shen and Ma (2019) for 11 experiments of this type), but it relies on the model describing the relationship between a target stimulus and the underlying probability of a correct or positive response rather than simply having the model describe the probability of response errors). With this goal in mind, we adapted the orientation memory task by Bae and Luck’s (2018) into an orientation perception task. The researchers originally used their task to investigate how well information held in working memory can be decoded from brain activity. What makes the task useful for the current study is that it allows participants to give a continuous response when asked to report the orientation of the target. Performance can then be quantified as the angular difference between the orientation of the target and the orientation reported by the participants, referred to as response errors. This continuous measure of response errors can utilize models such as the standard mixture model introduced by Zhang and Luck (2008) or the variable precision model by Fougnie and colleagues (2012) to quantify parameters of interest such as guess rate and precision. By extending this method to orientation perception, we can look at how target detection and perceptual variability are individually related to electrical brain activity during the task.

To address the question about the type of process underlying visual perception, we combined our adapted version of the visual orientation task with the standard mixture model and electroencephalography (EEG). Orientation estimation tasks are common in the visual perception literature (Fischer & Whitney, 2014) and the application of the standard mixture model to perception, and orientation perception, in particular, has been previously studied (Bays, 2016; Samaha et al., 2019). However, to our knowledge, this is a novel approach to quantify the effects of EEG brain activity on visual orientation perception using the standard mixture model to quantify performance. As a result, the purpose of the study was two-fold. First, we asked whether the standard mixture model is a good choice for quantifying orientation perception task performance by comparing the fits of other appropriate working memory models to the data. All the working memory models we tested made the same assumption that the distribution of errors could be separated in a uniform distribution representing guesses and a normal distribution of seen or remembered targets. The main difference between models was how the variability of the normal distribution got defined. This means that regardless of the model, there would be a standard deviation parameter (the standard deviation parameter could be defined by one value or two depending on the model). The uniform distribution quantified by a guess rate parameter may or may not be included depending on the model. Therefore, our second purpose was to test the relationship between EEG activity and the model parameters. We hypothesized that alpha activity prior to the target onset would be related to whether the target was later perceived or not which would be reflected as a modulation of the guess rate parameter or modulation of the mean SD/mode precision if there is no guess rate parameter. We also hypothesized that the precision of perceptual representations was a fixed ranged (*i.e*., based on the same distribution across trials) and that it would be related to post-target activity in the lower frequency ranges (4-7 Hz) which would be reflected as modulation of the standard deviation parameter. To address these questions and test our hypotheses, we modified an orientation memory task from Bae and Luck’s (2018) so that it probed participants’ perception of the target’s orientation rather than their ability to remember it. We then recorded EEG activity as participants performed the adapted task so we can see how brain activity varies with their perceptual performance.

## Materials and Methods

### Participants

Twenty-eight participants from the University of Alberta community participated in the study (age range = 17-35 years). Two participants were not included in the analysis due to excessive movement artifacts (more than 25% of trials rejected due to artifacts). Another two participants were excluded from the analysis due to having extreme outlying performance on the task (see Behavioral Analysis in the Results section for more details). Participants were all right-handed and had normal or corrected normal vision and no history of neurological problems. All participants gave informed written consent, were either compensated at a rate of $10/hr or given research credit for their time. The study adhered to the tenets of the Declaration of Helsinki and was approved by the Internal Ethics Board at the University of Alberta.

### Orientation Perception Task

Participants were seated 57 cm away from a 1920 x 1080 pixel^2^ ViewPixx/EEG LCD monitor (VPixx Technologies, Quebec, Canada) with a refresh rate of 120 Hz, simulating a CRT display with LED backlight rastering. The rastering, along with 8-bit digital TTL output triggers yoked to the onset and value of the top left pixel, allowed for submillisecond accuracy in pixel illumination times, which were confirmed with a photocell prior to the experiment. Stimuli were presented using a Windows 7 PC running MATLAB R2012b with the Psychophysics toolbox (Version 3; Brainard, 1997; Pelli, 1997). The code running the task was a modified version of the ColorWorkingMemoryExperiment.m code from MemToolbox (Suchow et al., 2013; memtoolbox.org). The modified version of color working memory experiment can be found here: https://github.com/APPLabUofA/OrientTask_paper/tree/master/OrientationTask. Video output was sent to the ViewPixx/EEG with an Asus Striker GTX760 (Fremont, CA) graphics processing unit.

Each trial began with a white fixation dot presented at the center of the monitor for 742, 783, 825, or 867 ms (target stimulus onset asynchrony; tSOA) after which the target appeared for 8.33 ms (one monitor refresh). The target was in the shape of a needle and was pointing toward one of 24 predefined evenly spaced directions so that all the orientations covered 360 degrees. The direction of the target was randomly selected on each trial. Of all the trials, 20% were randomly chosen not to have a target. All aspects of the target-present and target-absent trials were identical except that for the target-absent condition a blank interval replaced target presentation. A backward mask lasting for 8.33 ms with a constant 41.7 ms target-mask SOA (mSOA) appeared centrally. The mask was created by overlaying the target orientated in all 24 directions which created a star shape seen in Figure 1A. Following the mask offset, a 500 ms blank interval period occurred.

**Figure 1.**
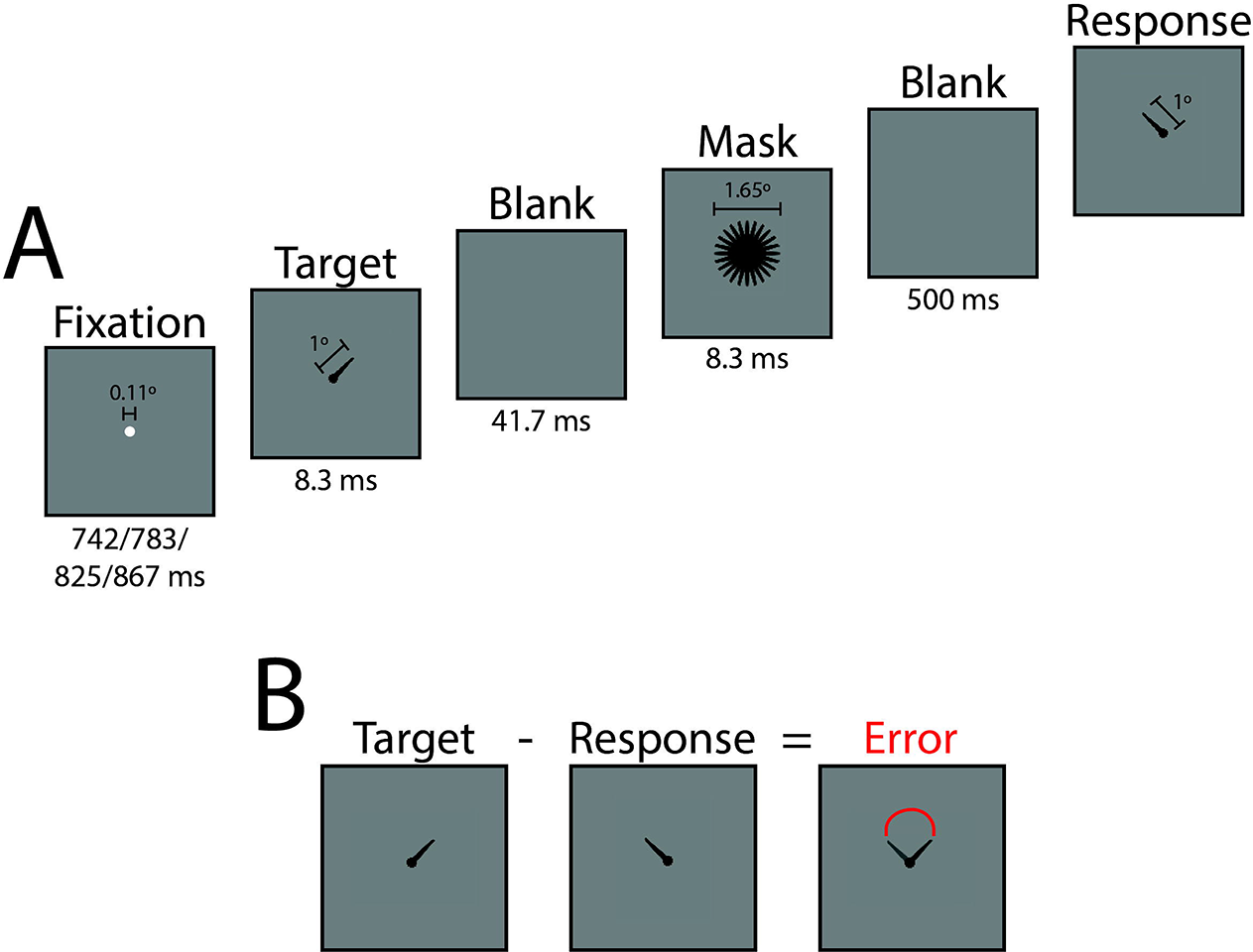
**A)** Sequence of task events with duration of each screen presentation and sizes of fixation, target, mask, and response stimuli. Sizes are in degrees of visual angle. **B)** Example of response error calculation. Response errors are reported in degrees.

After the blank interval, a response screen appeared with the needle in the center of the screen. Using the computer mouse, participants were asked to rotate the needle so that it was pointed in the same direction as the previous target. If participants detected a target but could not remember its orientation, they were asked to guess the orientation of the target. Participants could provide their response at their own pace. No feedback was given to participants. The next trial began immediately after a needle’s orientation was selected. See Figure 1A for a summary of the task sequence and the stimulus dimensions.

Participants completed seven blocks consisting of 48 trials each, along with 20 practice trials at the beginning of the experiment. Participants could rest at their own pace every 48 trials. Extensive written and verbal instructions were presented to participants prior to the practice trials. Instructions thoroughly explained and demonstrated each component that would compose a single trial.

Before the orientation perception task, participants performed a staircased target detection task that had the same parameters as the orientation perception task except that participants only reported whether they saw the target or not using the keyboard. The target color was a gray determined by a scalar value passed to the functions in Psychtoolbox. In the staircased target detection task, the target color value could range from the background color (making it not visible; corresponding value of 256) to black (making it the most visible; corresponding value of 0). This target gray value was adjusted throughout the task based on a 1-up/2-down staircasing procedure targeting a 0.6 target detection rate for each individual (Garcıra & Prins, 2016). The staircased task consisted of three blocks of 32 trials. The target gray value was determined for each participant by taking the average target gray value across the last two blocks of trials. These final average values ranged from 70 to 112 and were used as the target gray value in the orientation perception task.

The MATLAB code for the a staircased target detection task and the orientation perception task are available at https://osf.io/cw7ux/ and https://github.com/APPLabUofA/OrientTask_paper.

### EEG Recording

During the experiment, EEG data was recorded from each participant with a Brain-Amp 32-channel amplifier (BrainVision) using gelled low-impedance electrodes (actiCAP passive). Inter-electrode impedances were measured at the start of each experiment. All electrodes were arranged in the same 10-20 positions (Fp2, Fp1, F4, F3, F8, F7, FC2, FC1, FC6, FC5, C4, C3, CP2, CP1, CP6 CP5, P4, P3 P6, P5, P8, P7, PO4, PO3, O2, O1, Fz, FCz, Cz, Pz, and Oz). In addition to the 31 EEG sensors, a ground electrode was used, positioned at AFz. Two reference electrodes and the vertical and horizontal bipolar EOG were recorded from passive Ag/AgCl easycap disk electrodes affixed on the mastoids, above and below the left eye, and 1 cm lateral from the outer canthus of each eye. The bipolar channels were recorded using the AUX ports of the Brain-Amp amplifier. SuperVisc electrolyte gel and mild abrasion with a blunted syringe tip were used to lower impedances. Gel was applied and inter-electrode impedances were lowered to less than 5 kΩ for all electrode sites. EEG data was recorded online referenced to an electrode attached to the left mastoid. Offline, the data were re-referenced to the arithmetically derived average of the left and right mastoid electrodes.

Data were digitized at 1000 Hz with a resolution of 24 bits. Data were filtered with an online bandpass with cutoffs of 0.1 Hz and 250 Hz. The experiment was run in a dimly lit, sound and radio frequency-attenuated chamber from Electromedical Instruments, with copper mesh covering the window. The only electrical devices in the chamber were an amplifier, speakers, keyboard, mouse, and monitor. The monitor ran on DC power from outside the chamber, the keyboard and mouse were plugged into USB outside the chamber, and the speakers and amplifier were both powered from outside the chamber, and nothing was plugged into the internal power outlets. Any devices transmitting or receiving radio waves (e.g., cell phones) were removed from the chamber for the duration of the experiment.

### EEG Preprocessing

All analyses were completed using Matlab R2018b with the EEGLAB 13.6.5b (Delorme & Makeig, 2004) and CircStat (Berens, 2009) toolboxes, as well as custom scripts. After the data had been re-referenced offline, the bandpass FIR filter from EEGLAB was applied with lower and upper cut-offs of 0.1 Hz and 50 Hz. Data was segmented into 3000 ms epochs aligned to target onset (-1500 ms pre-target onset to 1500 ms post-target onset). The average voltage in the 200 ms baseline prior to the target was subtracted on each trial for every electrode, and trials with absolute voltage fluctuations on any channel greater than 1000 μ movements were then corrected with a regression-based procedure developed by Gratton, Coles, and Donchin (1983). After a second baseline subtraction with 200 ms pre-target, trials with remaining absolute voltage fluctuations on any channel greater than 500 μ further analysis. Data was then subjected to visual inspection and manual rejection of trials contaminated by artifacts. On average, 3% of trials were rejected during visual inspection. Other than the two participants mentioned earlier, none of the remaining participants had more than 20% of trials rejected in this manner.

### Data Analyses

Data analysis was performed using MATLAB R2018b (The MathWorks Inc, Natick, MA, USA) and EEGLAB 13.6.5b (Delorme & Makeig, 2004). All statistical analyses were conducted using MATLAB R2018b. Red-white-blue colormaps were created using the redblue.m function by Auton (2009) found here: https://www.mathworks.com/matlabcentral/fileexchange/25536-red-blue-colormap. The MATLAB code for data analysis is available at the GitHub repository https://github.com/APPLabUofA/OrientTask_paper and the raw data files are available at https://osf.io/cw7ux/.

#### Behavioral Data

Response errors on each trial were calculated by subtracting the orientation of the response stimulus, as reported by the participant, from the orientation of the target stimulus (see Figure 1B).

##### Comparing Model Fits

In addition to the standard mixture model proposed by Zhang and Luck (2008), the working memory literature has several other models similar to the standard mixture model but makes different assumptions about some of the parameters. Some of the working memory models are not appropriate for the current visual orientation perception task such as those that have an additional Von Mises distributions to account for “swapping” errors or errors where the participant report a distractor item rather than the target (Bays et al., 2009). On the other hand, the variable precision models were ones that could be appropriate for the current study. The original idea behind to model was that the precision of memory varies from trial-to-trial rather than being fixed as it is in the standard mixture model. This is done by Fougnie and colleagues (2012) by having the standard deviation parameter be distributed according to a higher-order distribution, we chose a Gaussian distribution in this case. The variable precision model proposed by van den Berg and colleagues (2012) has a precision (*i.e*., the inverse of variance) parameter drawn from a gamma distribution. . Neither paper presented clear justification for choosing one distribution over another, especially when the set size is always one, so we tested both distributions. In addition, we tested whether the variable precision models fit better to the response error data when they did not have the guess rate parameter compared to the standard mixture model and the variable precision models with a guess rate parameter. According to the variable precision models, what seems to be guessing is just low precision on that trial (Van Den Berg et al., 2012). If this were the case, the variable precision models without a guess rate parameter would fit the data better than the standard mixture models. On the other hand, the variable precision models do not, necessarily, preclude guessing. Fougnie et al. (2012) found that the models with a guess rate parameter described their data better than those without. To determine whether their findings extend to orientation perception data, variable precision models with a guess rate parameter were also tested.

We determined which model better fit the response errors using the model comparison routine in the MemToolbox (Suchow et al., 2013). We included the standard mixture model with the bias parameter in addition to the two variable precision models with and without the guess rate parameter. The goodness-of-fit measures used were the log likelihood and the Bayesian information criterion (BIC).

##### Standard Mixture Model

After determining the standard mixture model proposed by Zhang and Luck (2008) was the best fit to the current data set, the model was fit to each participant’s response errors using the maximum likelihood estimation routine in the MemToolbox (Suchow et al., 2013). According to the standard mixture model, response deviations from the actual target orientation reflect a mixture of trials where the target’s orientation was detected and trials where participants did not detect the target so guessed randomly. Therefore, the distribution of response errors consists of a mixture of a von Mises distribution (representing the trials where the target’s orientation was detected) and a uniform distribution (random guesses (*g*)). Parameter sigma (σ) is the standard deviation of the von Mises distribution, which represents the width of the response error distribution of trials that the target’s orientation was detected. Parameter *g* is the height of the uniform distribution representing the guessing probability. A third parameter, mu (μ), which is the mean of the von Mises distribution and represents systematic bias of the response error distribution was included in the standard mixture model of two participants because the Bayesian information criterion (BIC), calculated with the model comparison functions provided by MemToolbox (Suchow et al., 2013), indicated that the three-parameter standard mixture model provided better fits for those two participants (see Table 1). Although the three-parameter model was used for all analysis of those two participants, the systematic bias was much smaller than the spacing between adjacent target orientations (spacing was 15° whereas the two participants’ μ was -4.2° and 2.7°) indicating that those two participants had a slight clockwise and counterclockwise bias, respectively. The two-parameter standard mixture model was used for all analysis of the remaining 22 participants because it provided a better fit according to the BIC.

**Tabel 1.** List of goodness-of-fit measures comparing the standard mixture model to the standard mixture model with a bias parameter (µ) for each participant.

|  | Log Likelihood |  |  | Bayesian Information Criterion (BIC) |  |  |
| --- | --- | --- | --- | --- | --- | --- |
|  | Standard Model | Model + Bias | Difference | Standard Model | Model + Bias | Difference |
| Participant 1 | -1255.82 | -1255.33 | -0.49 | 2526.45 | 2532.88 | -6.44 |
| Participant 2 | -1213.93 | -1212.78 | -1.15 | 2442.77 | 2447.92 | -5.15 |
| Participant 3 | -1095.28 | -1094.58 | -0.70 | 2205.45 | 2211.50 | -6.05 |
| Participant 4 | -1093.29 | -1093.24 | -0.05 | 2201.25 | 2208.50 | -7.24 |
| Participant 5 | -1191.23 | -1190.92 | -0.31 | 2397.23 | 2404.01 | -6.77 |
| Participant 6 | -1113.21 | -1111.14 | -2.07 | 2241.28 | 2244.57 | -3.29 |
| Participant 7 | -965.54 | -964.92 | -0.62 | 1945.90 | 1952.06 | -6.16 |
| Participant 8 | -1022.30 | -1021.07 | -1.23 | 2059.48 | 2064.47 | -4.99 |
| Participant 9 | -1109.18 | -1109.12 | -0.06 | 2233.12 | 2240.39 | -7.27 |
| Participant 10 | -1422.57 | -1422.42 | -0.15 | 2859.94 | 2867.05 | -7.11 |
| Participant 11 | -1384.47 | -1384.46 | -0.00 | 2783.78 | 2791.20 | -7.42 |
| Participant 12 | -1017.58 | -1015.52 | -2.06 | 2049.58 | 2052.67 | -3.09 |
| Participant 13 | -1498.89 | -1498.55 | -0.34 | 3012.59 | 3019.32 | -6.73 |
| Participant 14 | -1071.57 | -1071.55 | -0.02 | 2157.98 | 2165.36 | -7.38 |
| Participant 15 | -1124.71 | -1123.84 | -0.87 | 2264.26 | 2269.94 | -5.68 |
| Participant 16 | -985.46 | -985.17 | -0.29 | 1985.36 | 1991.99 | -6.63 |
| Participant 17 | -1149.36 | -1149.26 | -0.11 | 2313.61 | 2320.84 | -7.23 |
| Participant 18 | -1136.58 | -1136.50 | -0.09 | 2287.86 | 2295.02 | -7.17 |
| Participant 19 | -1133.03 | -1132.91 | -0.12 | 2280.88 | 2288.04 | -7.16 |
| Participant 20 | -1276.89 | -1275.81 | -1.08 | 2568.44 | 2573.61 | -5.17 |
| Participant 21 | -1138.27 | -1133.04 | -5.23 | 2291.40 | 2288.37 | 3.03<sup>a</sup> |
| Participant 22 | -1003.69 | -988.80 | -14.89 | 2022.10 | 1999.69 | 22.41<sup>a</sup> |
| Participant 23 | -1011.18 | -1010.68 | -0.50 | 2037.17 | 2043.57 | -6.40 |
| Participant 24 | -1166.38 | -1162.88 | -3.49 | 2347.60 | 2348.04 | -0.44 |

*Note.* Difference is calculated as the goodness-of-fit measure of the standard mixture model with a bias model subtracted from the goodness-of-fit measure of the standard mixture model.
<sup>a</sup> Positive difference indicates the standard mixture model with a bias parameter ( $\mu$ ) fits better to the participants' response errors (*i.e.*, has a smaller BIC) than the standard mixture model.

#### Time-Frequency Analyses

To calculate the phase angle and power for each trial, we used Morlet wavelet transform of single trials using the newtimef() function of EEGLAB. A Morlet wavelet is a tapered sinusoid, constructed by modulating a sine wave with a Gaussian envelope. Wavelet transformation was created with 1.027 Hz linear steps and cycles starting from 2 cycles at 2 Hz to 12 cycles at 40 Hz. The output of this function was a matrix of complex values. The abs() function from MATLAB was used to get the instantaneous amplitude and the instantaneous phase angle of each trial was calculated using the angle() function from MATLAB.

EEG power data was converted to *Z* scores by applying a single-trial normalization procedure to the data from each participant at each electrode and frequency separately. This was done because it helps disentangle background from task-related dynamics, allows for comparison across different frequency bands and electrodes, and facilitates group-level analysis (Cohen, 2014). Each trial’s entire epoch (-700 ms to 800 ms relative to target onset) was used for the baseline during normalization because it has been shown to be robust to the effects of noisy trials (Grandchamp & Delorme, 2011). The downside to using the entire epoch for the baseline normalization is that sustained changes throughout the trial period become difficult to detect (Cohen, 2014). Ultimately, we considered this an acceptable trade-off for being able to compare effects in power and behavioral measures across participants without having to make as many assumptions about frequency bands and time-windows.

Another possible limitation of using the entire epoch for the baseline during normalization is that changes in pre-target power may be obscured by post-target activity. However, comparing effects across participants using raw EEG power is difficult due to individual differences caused by factors independent of the experimental manipulations (*e.g*., skull thickness) (Cohen, 2014). Therefore, when possible, the logarithmically transformed power was used. EEG power data was logarithmically transformed by applying a single-trial log10 transformation procedure to the data from each participant at each electrode and frequency separately. When the raw log transformed power is used, it is referred to as log power. EEG power that was converted to z-scores is called baseline normalized power.

#### Accurate vs Guess Trials

To test for significant differences in brain activity on trials where participants had small response errors compared to large response errors, we separated each participant’s trials based on the sigma value from the fits of their individual response errors to the standard mixture model.

This was done by defining each participant’s trials with response errors between -0.75σ and +0.75σ as “accurate,” and trials with response errors less than -1.5σ and greater than +1.5σ as “guesses” (Figure 2A). Trials where the participant clearly perceives the target are likely trials with a response error less than the participant’s overall response standard deviation, and trials the participant has little to no perception of the target are likely trials with a response error greater than just the participant’s response standard deviation. It should be noted that there are various reasons for participants to be accurate when they are guessing or have a large response error when they accurately perceived the target. However, such events are thought to be rare enough, or at least not systematic, that they will not unduly affect the overall distribution.

**Figure 2.**
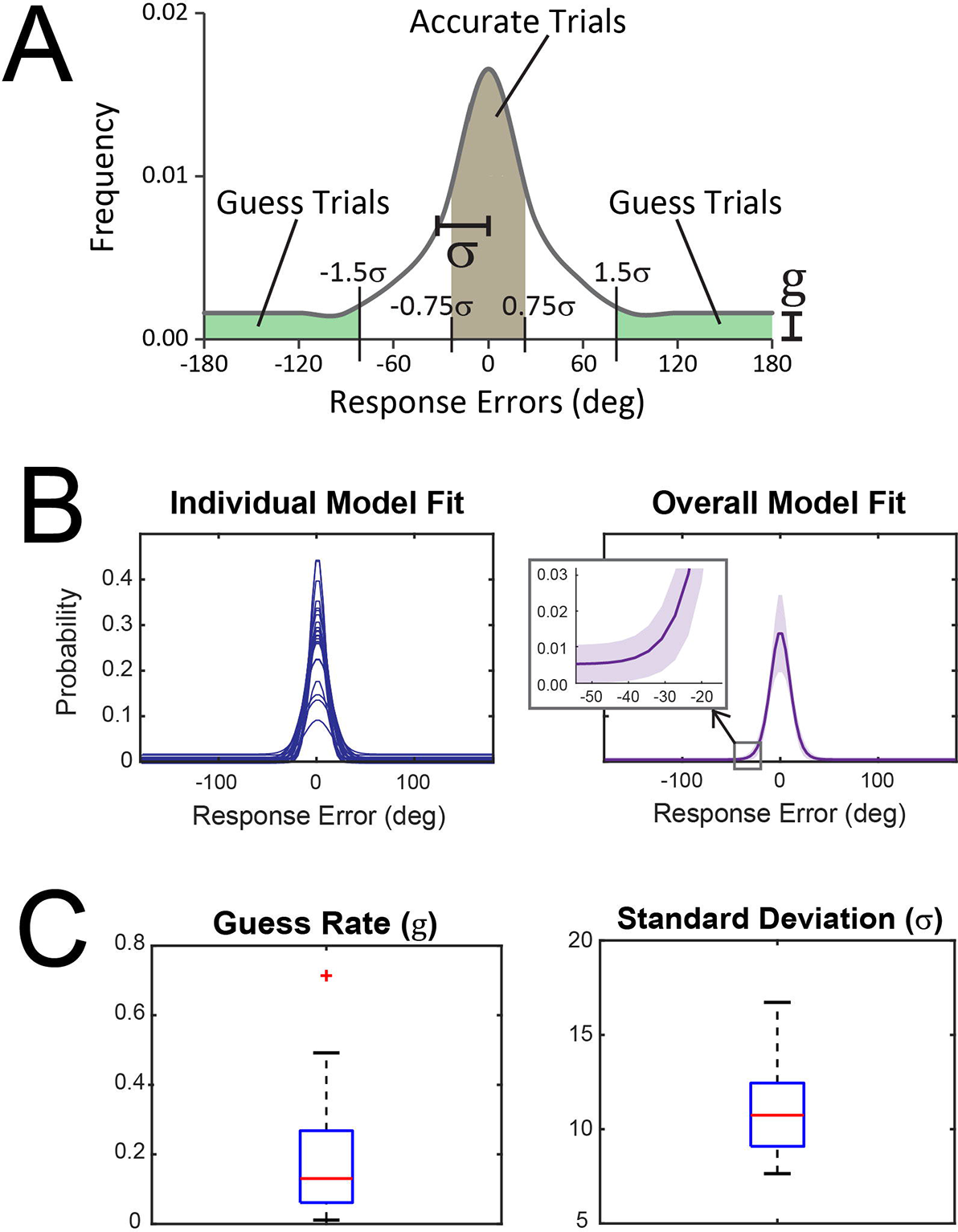
**A)** Representation of how trials were split into the two categories, Accurate and Guess. This was done by defining each participant’s trials with response errors between -0.75σ and +0.75σ as “accurate” (shaded brown region), and trials with response errors less than -1.5σ and greater than +1.5σ as “guesses” (shaded green regions). Trials that did not fall in either category were discarded from this analysis between the two trial categories. **B)** Left, model fit of each participant. Right, solid purple line is the average of participants’ model fit. Light purple area represents ±*SEM*. Zoomed in window shows upward shift of the averaged model fit showing a non-zero guess rate. **C)** Boxplots of parameter values (left) guess rate and (right) standard deviation estimated from model fits of participants’ response errors.

The main reason we separated trials into “accurate” or “guess” was to see if the standard mixture model parameters could be used to categorize trials in a meaningful way. Though this method resulted in excluding some trials, namely ones that fell between the cutoffs for accurate and guess, the outcome provided insight into how brain activity differs between different levels of objective perceptual performance.

##### ERP Analyses

To remove the activity elicited by the mask without removing activity resulting from the interaction of mask and target, the catch trials (mask-only) average was subtracted from the orientation detection trials (target-plus-mask) average. ERP data was submitted to a repeated-measures, two-tailed permutation test based on the *tmax* statistic (Blair & Karniski, 1993) using the mxt_perm1() function from the Mass Univariate ERP Toolbox (Groppe et al., 2011). The time windows of interest were the P1 (80-140 ms), N1 (140-200 ms), P2 (200- 255 ms), N2 (255-360 ms), and P3 (360-500 ms) components. The ERP component time windows were selected based on previous literature (Koivisto & Revonsuo, 2003, 2010). All 31 brain electrodes were included in the test. 100,000 random within-participant permutations were used to estimate the distribution of the null hypothesis and the familywise alpha (α) was set to 0.05. Based on this estimate, critical *t*-scores of +/-3.68 (*df* = 23) were derived. Any t-scores that exceeded the critical *t*-score were considered statistically significant.

##### Baseline Normalized EEG Power Analysis

To analyze differences in EEG baseline normalized power between guess and accurate trials, nonparametric permutation testing with a pixel-based multiple-comparison correction procedure (Cohen, 2014) was used to analyze differences in EEG band power between guess and accurate trials. The pixel-based multiple-comparison correction method involves creating one distribution of the largest positive pixel value and another distribution of the largest negative pixel value from each iteration of the permutation testing. After all iterations, the statistical threshold is defined as the value corresponding to the 2.5^th^ percentile of the smallest values and the value corresponding to the 97.5^th^ percentile of the largest values which are the thresholds corresponding to an α of 0.05. Any pixel that has a value exceeding the upper or lower value is considered significant. The pixel-based method corrects for multiple comparisons by creating two distributions based on map-level information instead of pixel-level information. In other words, this method results in two distributions of the most extreme null-hypothesis test statistical values across all pixels rather than calculating null-hypothesis distributions for each pixel (see Cohen (2014) for further details about pixel-based multiple-comparison correction method). All analysis using nonparametric permutation testing with pixel-based multiple-comparison correction performed 10,000 iterations per test. To obtain more stable estimates from permutation testing, we ran a “meta-permutation test” by repeating the pixel-level permutation procedure 10 times and then averaging the results (Cohen, 2014). It needs to be pointed out that a “significant effect” determined by pixel-based permutation testing should not be considered a precise estimate in the temporal and frequency domains. Although pixel-based permutation testing is more stringent than cluster-based permutation tests (Cohen, 2014), caution should still be used when interpreting “significant” differences, especially if the temporal and frequency range of each pixel is relatively small.

##### EEG Phase Analysis

To determine whether the mean phase values significantly differ between accurate and guess trials, we used the circular Watson–Williams (W-W) test which was calculated using the PhaseOpposition.m function by VanRullen (2016). We chose the parametric circular W-W test because it has shown to be equivalent to the non-parametric phase opposition sum (POS) measures under most conditions and performed better in situations where either the relative trial number or the ERP amplitude differed between the two trial groups (VanRullen, 2016). The statistical significance of the W-W test across participants was determined by combining the individual-level *p*-value time-frequency map at each electrode across participants using Stouffer’s method (Stouffer et al., 1949; VanRullen, 2016), which transforms individual *p*- values into *z*-scores, combines them across participants, and converts the resulting *z*-score to a combined probability. *P*-values were then corrected for multiple comparisons across time points and frequencies at each electrode using the false discovery rate (FDR) procedure described in Benjamini and Yekutieli (2001). Effects that satisfied a 5% FDR criterion were considered significant.

#### Single-Trial EEG Activity and Response Errors

To test the correlation between time-frequency log power and degree of response error, Spearman’s rho (*r_s_*) correlation coefficients were calculated using a nonparametric permutation testing approach with the pixel-based multiple-comparison correction procedure described above. The null-hypothesis distribution was created by shuffling power values and response errors on each trial with respect to each other. This provided a data-driven test of the null hypothesis that there is no consistent relationship between degree of response error and EEG power.

To look at whether task performance is related to oscillatory phase, and if yes, at what frequency, we used the weighted inter-trial phase clustering (wITPC) (Cohen, 2014; Cohen & Voytek, 2013). The logic behind the inter-trial phase coherence (ITPC) is that a systematic relation between EEG phase and behavioral outcome should result in a higher-than-chance ITPC in each of the trial subgroups. However, if the phase of the EEG signal is randomized and unpredictable, the distribution of phases at a given time period should follow a uniform distribution over all trials. The problem with ITPC is that it assumes EEG phase is relevant to experimental measures only when phase values are similar across trials (van Diepen & Mazaheri, 2018). Unlike ITPC, wITPC is sensitive to modulations of phase values even if those phases are randomly distributed across trials as would be expected if response errors (which differs from trial to trial) were modulated by oscillatory phase (Cohen, 2014; Cohen & Voytek, 2013).

The wITPC was computed for each participant as the resultant vector length, or ITPC, of phase angles across trials once the length of each vector has been weighted by a variable of interest (in this case, each trial’s phase vector is weighted by the degree of response error on that trial; see Figure 6A for example of computation) (Cohen, 2014; Cohen & Voytek, 2013). For statistical testing, a null-hypothesis distribution was created by shuffling the phase values relative to trial response error 10,000 times (see Figure 6A middle and bottom left). The wITPCz was calculated as the wITPC standardized relative to the null-hypothesis distribution, providing a *z*-value corresponding to the probability of finding the observed response error–phase modulation by chance, given the measured data. As was done for the parametric circular W-W test, statistical significance of the wITPCz across participants was evaluated by combining the individual-level *p*-value, calculated from the *z*-values, time-frequency map at each electrode across participants using Stouffer’s method (Stouffer et al., 1949; VanRullen, 2016). *P*-values were then corrected for multiple comparisons across time points and frequencies at each electrode using the false discovery rate (FDR) procedure described in Benjamini and Yekutieli (2001). Effects that satisfied a 5% FDR criterion were considered significant.

We chose to use the phase opposition measure and the wITPCz even though they are both quantifying phase coherence because they provide slightly different but complementary information about the effects of phase. The phase opposition measure provides insight into whether there is an overall consistent difference in mean phase values between trials separated by model parameter defined categories (*i.e*., “accurate” and “guess”). On the other hand, the weighted single-trial phase modulation metric (*i.e*., wITPCz) provides information about response error-specific modulations of phase values irrespective of the model. In other words, the circular W-W test reflects differences between the mean phase of the binned trials while wITPCz reflects differences in phase as it relates to the continuous measure of response errors. Also, the wITPCz does not rely on phase values being consistent over trials, they only need to be consistently related to response errors (Cohen & Cavanagh, 2011). This is an important distinction when trying to determine how much of the phase modulation is an artifact of the stimulus-evoked activity and how much is related to the difference in task performance.

#### Relationship Between Log Power and Standard Mixed Model Parameters

To determine how mixture model parameters standard deviation and guess rate varied as power varied, a median split of trials according to raw power at each time and frequency point was done for each participant at each electrode separately. EEG power from the Morlet Wavelet transformation was logarithmically transformed by applying a single-trial log10 transformation then trials were split by whether they were above or below the median power at each time point and frequency. This was done separately for each participant at each electrode. The standard mixture model was then fit to each set of response errors on the high power and low power trials to get model parameter values. This meant that every time-frequency point had a standard deviation and guess rate from trials with high power and low power. The “high power” and “low power” parameters were then averaged across a frequency band (2-3 Hz, 4-7 Hz, 8-14 Hz, 15-29 Hz or 30-40 Hz) and then tested statistically with the same procedure as the ERP analysis except each time point was tested rather than averaging across a time window and the alpha level was set to 0.01 to control for the familywise error rate (Bonferroni corrected alpha level α_corr_ = 0.05/5 to account for the five frequency bands). Figure 3 gives a visual overview of each step in this analysis.

**Figure 3.**
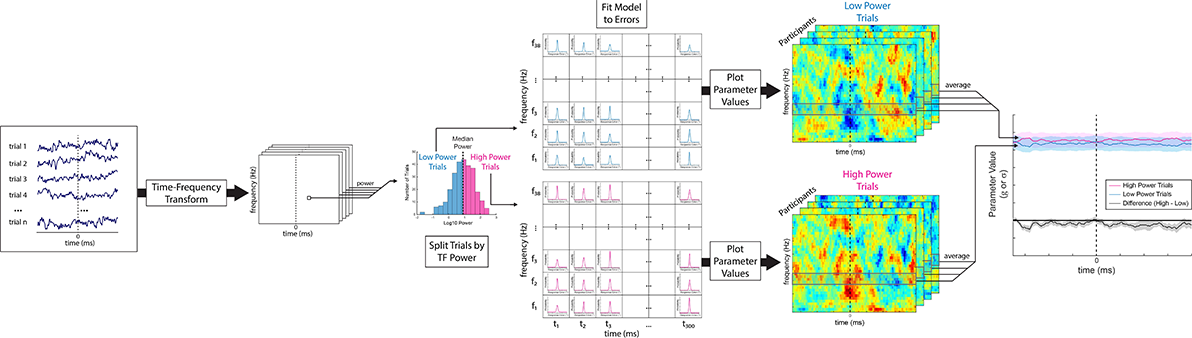
Illustration of the analysis looking at how the estimated parameter values from the standard mixture model varied across an epoch when trials were separated by the power within a frequency band at each time point. First, time-frequency transformation (e.g., Morlet wavelet transformation) of single-trial data was used to calculate raw power of every time and frequency point for each trial. Trials were split by median power at each time and frequency point then the standard mixture model was fit to the trials’ response errors. This was done for each participant and electrode separately. An average of the parameter values (*i.e.,* standard deviation and guess rate) from the “high power” and “low power” trials were calculated across the five frequency bands (2-3 Hz, 4-7 Hz, 8-14 Hz, 15-29 Hz or 30-40 Hz) at each time point and then were compared statistically using a repeated measures, two-tailed permutation test based on the *tmax* statistic (Blair & Karniski, 1993).

Considering the timing and frequency of these significant effects, it is likely they reflect the same processes measured by the ERP components. To test this idea, a procedure like the one described above was applied to the ERP data so that trials were split by the average amplitude of each ERP component rather than at each time point. The “high amplitude” and “low amplitude” fitted model parameters were tested statistically with the same procedure as the accurate vs guess ERP analysis.

For comparison with the ERP results, the guess rate and standard deviation parameters from high and low 2-3 Hz and 4-7 Hz log power trials were averaged across the time windows used for each ERP component: P1 80-140 ms (P1), 140-200 ms (N1), 200-255 ms (P2), 255-360 ms (N2), and 360-500 ms (P3). These were submitted to the same statistical procedure as the ERP components except the alpha level was Bonferroni corrected to 0.025 to account for testing two frequency bands. The two frequency bands were chosen because they are the only ones that have shown significant effects across all previous analyses.

Stepwise multiple regression analyses were performed for the standard mixture model parameters. Details about the methods and results can be found in Supporting Information.

## Results

### Comparing Model Fits

We tested the best fitting model using the goodness-of-fit measures log likelihood and the Bayesian information criterion (BIC). The results are presented in Figure 4 and the *mean* ± *SEM* of the fitted parameters for all the tested models are in Table S1. Overall, the BIC indicates the standard mixture model fits the data better than any of the variable precision models and the standard mixture model with a bias parameter. As Fougnie and colleagues (2012) found, the models also performed better when they included a guess rate parameter. The log likelihood indicates the standard mixture model with a bias parameter is better than the standard mixture model and the variable precision models. The differences between the BIC and log likelihood metrics can be attributed to the BIC having a penalty for model complexity whereas the log likelihood does not control for those factors.

**Figure 4.**
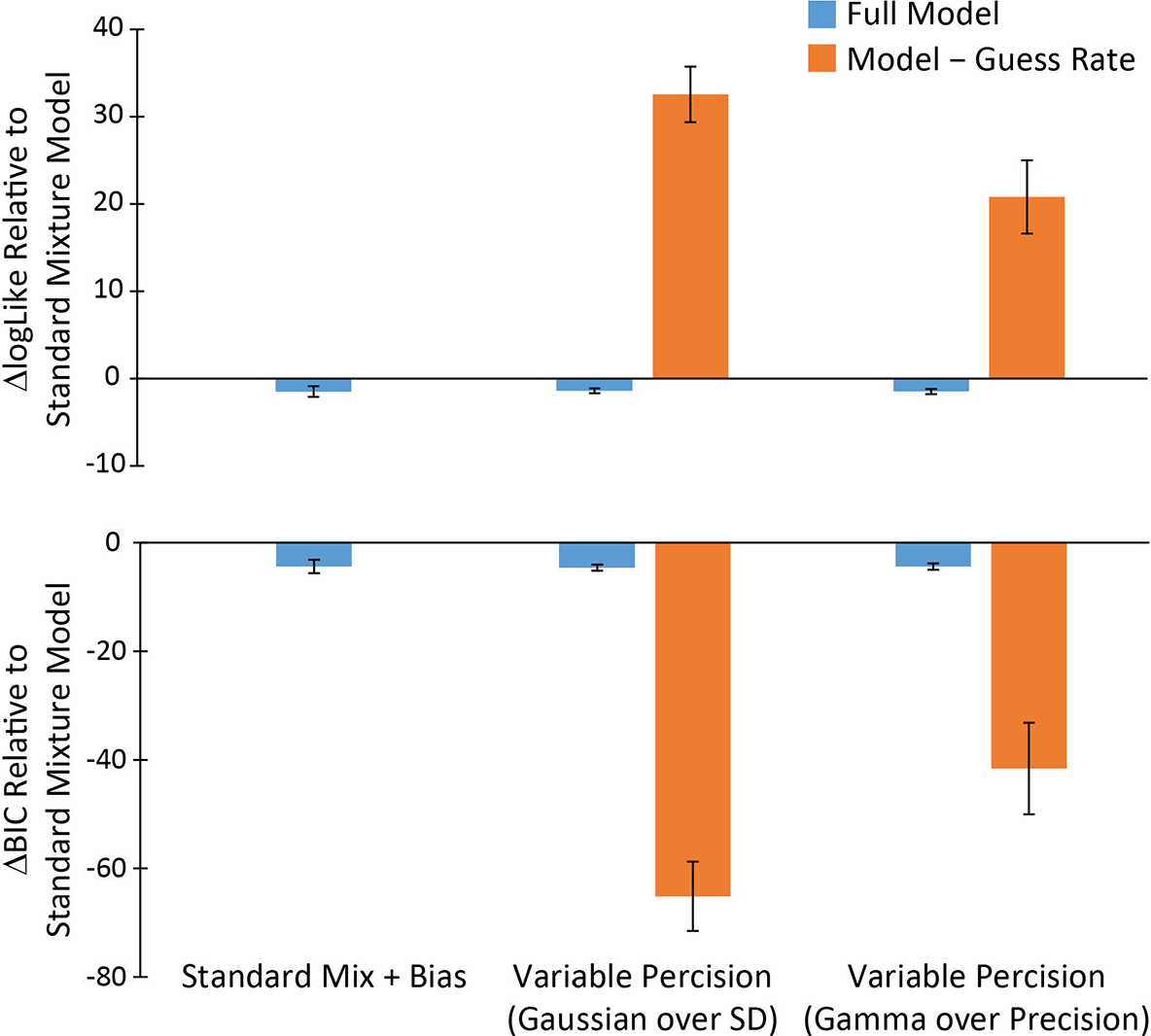
Goodness-of-fit measures relative to the standard mixture model. The models compared to the standard mixture model were the standard mixture model with a bias parameter and the two variable precision models with and without a guess rate parameter (Model – Guess Rate). Top, the difference in log likelihood values compared to the standard mixture model with positive values favoring the standard mixture model. Bottom, the difference in Bayesian Information Criterion (BIC) values compared to the standard mixture model with negative values favoring the standard mixture model. Log likelihood favors the standard mixture model with a bias parameter (µ) over the standard mixture model and both variable precision models. The BIC favors the standard mixture model over the other three models. Plots are the means and error bars are ±*SEM*.

### Accurate vs Guess Trials

Two participants were excluded before further analysis because one had a guess rate more than three *IQR*s from the median and the other had a guess rate of 3.9e-15 indicating that the staircasing procedure did not work properly for this individual. These participants were not included in further analysis. Figure 2B shows the fit of the standard mixture model (or standard **mixture** model with a bias parameter for the participants mentioned in the Methods section) to each participant’s response errors as well as the average fit of response errors across participants. The remaining 24 participants had a mean guess rate of 0.19 (*SD* = 0.18) and mean standard deviation (σ) parameter of 11.1 (*SD* = 2.5). The boxplots in Figure 2C summarizes the distributions.

### ERP Analysis

The ERPs from accurate and guess trials (Figure 5A) showed no statistical difference for the first 200 ms following target onset across all electrodes. A divergence in the waveforms can be seen in the P2 (200-300 ms) component for the frontal, central, and centroparietal electrodes (Figure 5B left) in that the voltage of the guess trial ERPs was much attenuated compared to the accurate trial ERPs. On the other hand, the voltage of the N2 (255-360 ms) component was more negative in the guess trials than accurate trials (Figure 5A) and this difference was only significant in the right frontocentral, central, centroparietal and parietal electrodes (Figure 5B middle). Finally, the P3 (360-500 ms) component from the guess trials had a similar attenuation as was seen in the P2 component (Figure 5A), but the distribution was more posterior with significant effects seen in the right central, centroparietal and parietal electrodes as well as bilateral parietooccipital and occipital electrodes (Figure 5B right).

**Figure 5.**
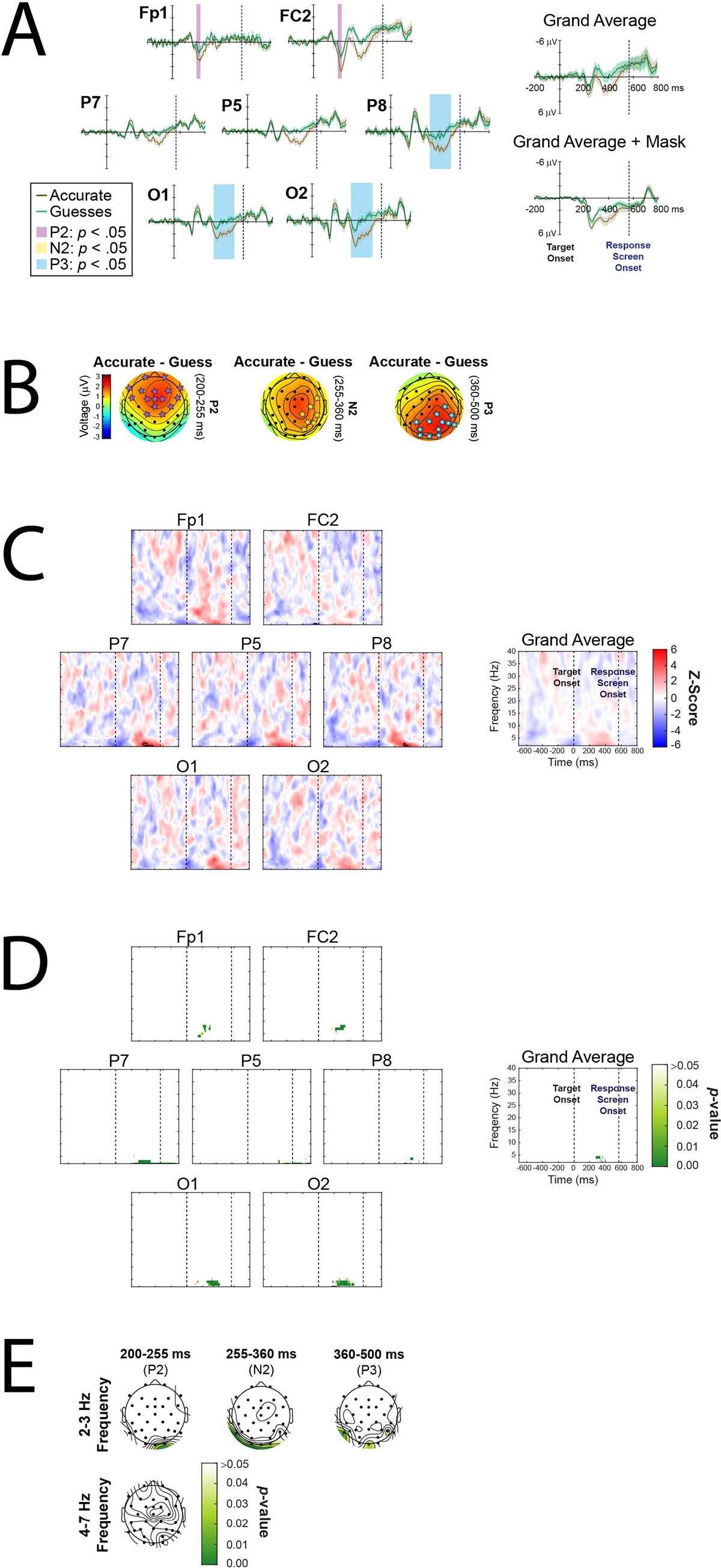
**A)** ERPs of accurate trials and guess trials aligned to target onset at selected electrodes and averaged across all electrodes (Grand Average). Light brown and light green shaded areas around waveforms represent ±*SEM* of accurate trials and guess trials, respectively. Mask trial EPRs have been subtracted out of the ERPs to remove mask-related brain activity except for the Grand Average + Mask ERPs plot. Time period shaded in purple indicates significant difference between accurate and guess trials 200-255 ms (P2) post-target. Blue shaded time period indicates significant difference between accurate and guess trials 360-500 ms (P3) post-target. **B)** Topographies of the voltage distribution difference between accurate and guess trials. The time periods in each topography are as follows: 200-255 ms (P2), 255-360 ms (N2), and 360-500 ms (P3) post-target. Stars indicate electrodes with significant differences between accurate and guess ERPs. Mask trial activity has been subtracted out of the ERPs to remove mask-related brain activity. **C)** Analysis results from comparing the baseline normalized power of accurate vs guess trials. Time-frequency plots showing the results of the statistical analysis of the difference in power between accurate vs guess trials at each time-frequency point at selected electrodes and the average of all electrodes (Grand Average). Black contour denotes statistically significant differences (after applying the pixel-based multiple-comparison correction procedure) at *p* < .05. **D)** Phase analysis of accurate vs guess trials. Time-frequency plots showing the *p*-values (after applying FDR correction) from the statistical analysis testing differences in the mean phase of accurate trials vs guess trials. Plots are from selected electrodes and the average of all electrodes (Grand Average). Time-frequency points with *p*-values above the threshold of FDR correction for multiple comparisons were set to 1. **E)** Topographies of the *p*-value distribution averaged across the 2-3 Hz frequency (top) and 4-7 Hz frequency (bottom) indicating significant differences in the mean phase of accurate trials vs guess trials. Time-frequency points with *p*-values above the threshold of FDR correction for multiple comparisons were set to 1. The time periods in each topography are the same as used for the ERP analysis: 200-255 ms (P2), 255-360 ms (N2), and 360-500 ms (P3) post-target. Only the time periods and frequency bands that had significant effects after averaging over the time window are shown.

### EEG Power Analysis

Pixel-based permutation test indicated significant differences between accurate and guess trials within the 2-4 Hz frequency range which was observed to start around 310 ms post-target onset in P8 with a duration of about 70 ms and 350 ms post-target onset in P7 with a duration of around 120 ms (Figure 5C).

Overall, there was a trend for increased 4-7 Hz power in accurate trials compared to guess trials, especially in left frontal and right parietal areas, though this difference was not significant. This lack of significance might be due to too much variability in when the 4-7 Hz power changed during the trial. It is also possible that 4-7 Hz activity reflects a perceptual process that could occur in both accurate and guess trials, though more often or to a greater degree in one type of trial compared to the other.

Finally, there were a brief period (20-30 ms) where guess trials had significantly more baseline normalized power than accurate trials at around 2 Hz right before target onset in FC2 (around -50 ms; Figure 5C) and Cz (around -70 ms; not shown). However, because of the wavelet parameters used and the timing being around a large evoked response, the timing of the difference is likely smeared backward so that the effect probably occurred after the target had been presented (Brüers & VanRullen, 2017; Herrmann et al., 2014; VanRullen, 2011). It should be noted that no other analysis yielded a significant effect immediately before or after target onset suggesting that these results are false positives, or the other analyses lacks the power to detect the effect. The very short duration of an effect at such a low frequency suggests the former is more likely. If it is the latter, the timing suggests it might have something to do with the mask onset (*e.g*., anticipation of the mask stimuli) rather than the target. However, the current study was not designed to investigate the masking stimulus making it difficult to determine the truth behind the observed effect.

### EEG Phase Analysis

Similar to the baseline normalized power results, significant differences in mean phase between accurate and guess trials were found in the 2-7 Hz frequency ranges following target onset (Figure 5D). Central and bilateral frontal, frontocentral, and central electrodes show significant phase differences in 4-7 Hz starting between 150-200 ms and terminating before 350 ms post-target (not shown). The duration of this effect varies so that the more central electrodes tended to have longer durations than those placed more laterally on the head. This effect was more prevalent in the right hemisphere in the early period.

The most lateral parietal and centroparietal electrodes on the left side of the head show significant phase differences within 2-3 Hz after the response screen onset (Figure 5E). On the other hand, Pz, P3 (Figure 5D), P4, and PO3 electrodes had no significant effects in phase while P6 and P8 had brief periods of significant phase differences starting a little after 300 ms until 500 ms post-target in the 2-3 Hz frequency range. P7 showed a similar significant difference in mean phase at 2 Hz starting around 225 ms and continuing until more than 200 ms after the response screen had been presented (about 770 ms post-target; Figure 5D). The occipital electrodes had significant phase effects primarily in the 3-5 Hz range starting a little after 100 ms on the left (not shown) and 150 ms on the right but were brief time periods until 200 ms post-target which then had significant differences lasting for about 200 ms (Figure 5D). PO4 electrode showed a similar difference, but the effect was less continuous and had larger *p*-values (*i.e*., smaller phase difference between accurate and guess trials) than the occipital electrodes; however, PO4 also had significant differences within 2-3 Hz frequency between 400 and 500 ms (Figure 5D).

It should be noted that the time window of significant phase opposition overlaps with the significant differences in ERP amplitudes. ERP amplitudes have been shown to have a “masking” effect on phase opposition measures resulting in a decrease in their statistical power (VanRullen, 2016) while at the same time the stimulus-evoked activity results in temporal distortion of oscillatory activity towards earlier latencies (Brüers & VanRullen, 2017). While the circular W-W test is relatively robust against the detrimental effect of ERPs, it does not negate their influence entirely (VanRullen, 2016). Therefore, it is important to be aware that the significant effects of phase are affected by the stimulus evoked activity and the different ERP amplitudes between accurate and guess trials.

### Single-Trial EEG Activity and Response Errors

To see whether response errors were related to trial-by-trial changes in pre-target alpha power, the Spearman’s rho correlations were calculated between log power and degree of response error on each trial. Our results did not allow us to reject the null hypothesis. Follow-up analysis on the entire time-frequency space also yielded no significant results (not shown).

To examine the relationship between response errors and the distribution of phase values, we used the wITPCz. As can be seen in Figure 6B, phase was primarily modulated by degree of response errors in the 2-3 Hz frequency range in the posterior electrodes starting at around 200 ms post-target and lasting until response screen onset. PO3 was similar except the effect started at around 250 ms (not shown), but PO4 showed an effect starting at 350 ms and lasting until almost 100 ms after the response screen onset (not shown). Parietal electrodes show a significant effect in the low frequency bands starting at around 200 ms until about 800 ms post-target (Figure 6B, top and bottom rows). Centroparietal and central electrodes had a significant relationship between phase at 2-3 Hz and response errors at around 300-400 ms post-target and lasted until after response screen onset. All frontocentral electrodes had response errors significantly related to 2-3 Hz phase starting between 200 ms and 300 ms post-target and lasting until shortly after response screen onset. Out of all the frontal electrodes, only Fz and F3 had a significant 2-7 Hz phase relationship to response errors (not shown).

**Figure 6.**
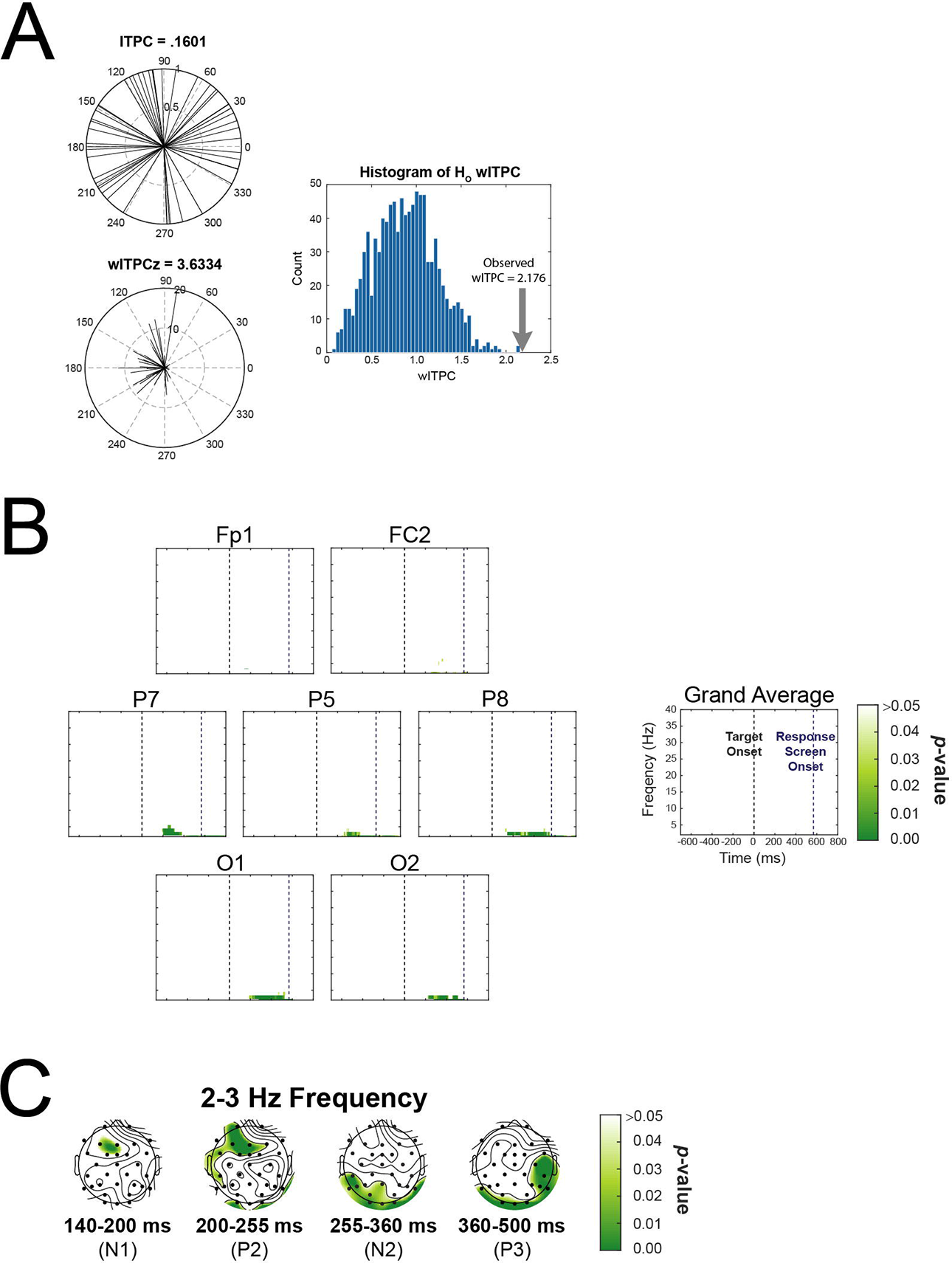
**A)** Example computation of weighted inter-trial phase clustering (wITPC) to relate single-trial phase to degree of response error. Top left, single-trial prestimulus phase vectors are shown as black lines and are not clustered across trials due to the randomization of the fixation length, leading to a low resultant vector length (i.e., low ITPC). Right, histogram of null-hypothesis vector lengths created by shuffling the mapping of the trial response error values to the trial phase values. The observed vector length (large arrow) is calculated from the distribution shown in bottom left panel. Bottom left shows how the length each trial’s phase vector is scaled by that trial’s degree response error and a weighted ITPC is computed, reflecting the relationship between the distribution of phase angles and degree of response error even though the distribution of the phase angles themselves is uniform (as seen in the top left plot). The wITPCz value is the wITPC standardized relative to the null-hypothesis distribution seen in the histogram plot. Example based on figure by Cohen (2014). **B)** Time-frequency plots of analysis relating single-trial phase activity and response errors. Significant *p*-values indicating that the normalized distance of the observed wITPC (i.e., wITPCz) is significantly different from the distribution of null hypothesis wITPC values. This measure represents the relationship between the distribution of phase angles and the degree of response error on each trial. Plots are only of selected electrodes and the average of all electrodes (Grand Average). Time-frequency points with *p*- values at or above .05 were set to 1. **C)** Topographies of the *p*-value distribution (after applying FDR correction) averaged across the 2-3 Hz frequency indicating that the normalized distance of the observed wITPC (i.e., wITPCz) is significantly different from the distribution of null hypothesis wITPC values. This measure represents the relationship between the distribution of phase angles and the degree of response error on each trial. Time-frequency points with *p*-values above the threshold of FDR correction for multiple comparisons were set to 1. The time periods in each topography are the same as used for the ERP analysis: 120-200 ms (N1), 200-255 ms (P2), 255-360 ms (N2), and 360-500 ms (P3) post-target. Only showing time periods and frequency bands that had significant effects after averaging over the time window.

### Relationship Between Log Power and Standard Mixed Model Parameters

A repeated measure, two-tailed permutation test indicated that significant differences in parameter values between high and low power trials were within the 2-3 Hz frequency band following target onset. Figure 7A shows the electrodes that had significant differences in guess rate and Figure 7B shows the electrodes that significant differences in the standard deviation (σ) parameter. No other electrodes had significant parameter value differences. Most of the effects are seen in the occipital and parietal electrodes except for the left frontal electrode which had a significant difference in guess rate at the later time points than observed in the other electrodes. The left parietal and occipital electrodes showing significant effects in standard deviation were not the same electrodes showing significant effects in guess rate. Interestingly, both the guess rate and standard deviation were higher in trials with low 2-3 Hz log power than high. No significant differences were seen in the 2-3 Hz frequency band prior to 250 ms post-target onset and did not occur later than 100 ms before the response screen appeared. It should be noted that, unlike the accurate vs. guess analysis, these results are based on the log power so the lack of pre- target effects cannot be an artifact of the normalization process.

**Figure 7.**
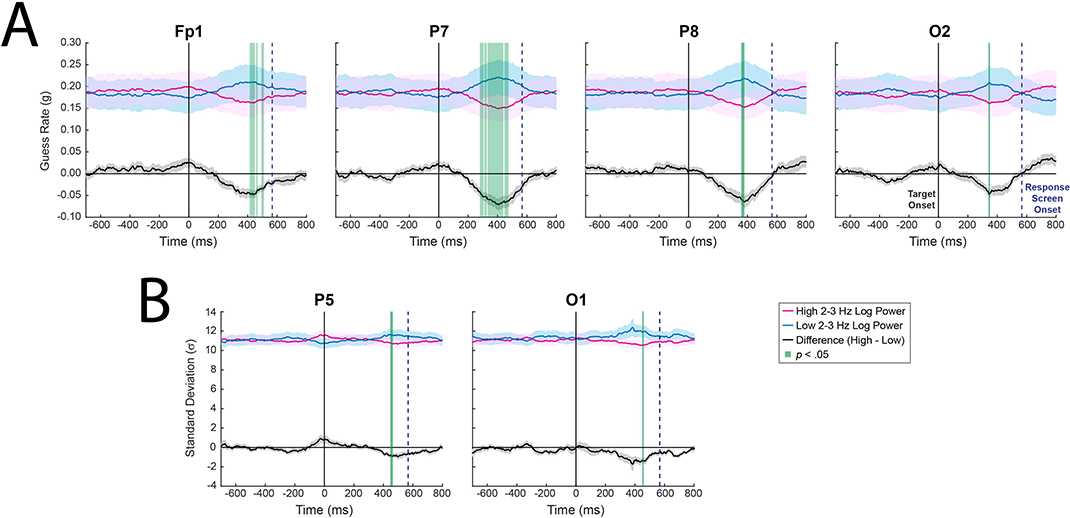
A) Plots of the fluctuations in the guess rate parameter across time in trials with high and low log power in the 2-3 Hz frequency band. No other frequency band had significant differences in guess rate between high and low power trials. Only electrodes with significant effects are shown. Green shaded regions indicate time points where guess rate in high and low power trials significantly differed. Shaded regions around waveforms are ±*SEM*. **B)** Plots of the fluctuations n the standard deviation (σ) parameter across time in trials with high and low log power in the 2-3 Hz frequency band. No other frequency band had significant differences in standard deviation between high and low power trials. Only electrodes with significant effects are shown. Green shaded regions indicate time points where the standard deviation parameter in high and low power trials significantly differed. Shaded regions around waveforms are ±*SEM*.

When the parameter values on high and low log power trials were averaged over the ERP time windows, a similar pattern of effects were seen in the 2-3 Hz frequency band. There was an overall trend for higher guess rates (Figure 8A) and larger standard deviations (Figure 8B) on trials with lower log power. The most notable difference was a significant difference in guess rate on trials with high and low 4-7 Hz log power at Pz in the 200-255 ms time window. Interestingly, when compared to the guess rates on trials with high and low P2 amplitudes, the ERP component during that time period, the significant differences are concentrated in the frontal and central electrodes. Based on visual inspection, the 4-7 Hz log power seems to contribute to P2 differences, but only weakly.

**Figure 8.**
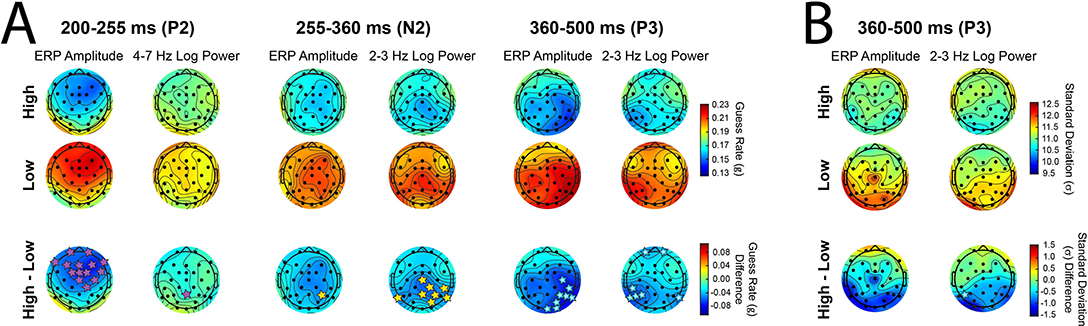
PERCEPTUAL QUALITY AS A FUNCTION OF EEG ACTIVITY 54 Topographies of the **A**) guess rate (g) and **B**) standard deviation (σ) parameters 1145 from fitting the standard mixture model to trials categorized as having high (top row) or low (middle row) ERP amplitude compared to the trials categorized as having high (top row) or low (middle row) log power in the specified frequency band. The bottom row shows the differences in the parameter values between high and low trials. The time period in each topography are the windows over which the mean amplitude was calculated for the ERP analysis and the windows over which the guess rate (g) or standard deviation (σ) parameters were averaged for the log power analysis (see igure 3 for more details). All time periods are relative to target onset. On the bottom row, stars indicate electrodes with significant differences between parameter values. The alpha level for the ERP amplitude and log power statistical comparisons was 0.05 and 0.025, respectively. Only the time periods and frequency bands with significant effects are shown except for the standard deviation (σ) parameters from fitting the ERP amplitude (**B**, left column) which were included for comparison.

In contrast, the significant effects for the 2-3 Hz frequency band activity during the N2 ERP (255-360 ms) is obscured for the ERP component itself. Only P6 had a significant difference in guess rate based on differences in N2 amplitude where most right centroparietal, parietal, and parietooccipital electrodes exhibited significant guess rate differences as well as the left parietal electrode P7.

Finally, the large P3 ERP showed a significant difference in guess rates at the more posterior electrodes though these differences were primarily on the right side. In comparison, during the P3 time window (360-500 ms) the more lateral parietal and centroparietal electrodes on the left side and the most lateral parietal electrode on the right side (P8) showed a different difference in guess rate on trials with high vs low 2-3 Hz log power. Interestingly, Fp1 and Fp2 also showed a significant difference in guess rates at 2-3 Hz frequency. Consistent with the time courses shown in Figure 7B, it was only during this late time period that a significant difference was seen in the standard deviation (σ) parameter values. Like the guess rate parameter, trials with high 2-3 Hz log power had greater standard deviations (σ) significant difference was only observed at the left posterior electrode, P5, though there was a trend for the bilateral occipital and parietal electrodes to have a relatively large standard deviation σ) on the low 2-3 Hz power trials (Figure 8B, middle row). In comparison, there were no ERP amplitudes showing a significant difference in the standard deviation (σ) parameter.

## Discussion

There were two goals for the present study. First, we asked what working memory model best fit to continuous response measures on an orientation perception task? The second goal of the study was to ask what might the relationship be between the parameters of the best-fitting model and EEG activity?

### Comparing Model Fits

Since perceiving the visual stimuli usually precedes remembering those same stimuli, some of the assumptions built into the visual working memory models are applicable to visual perception. Namely, the assumption that there are a set of targets that are remembered and a set of targets that are not remembered and they can be represented by two distributions which would translate to the current task as a set of targets that are seen and another set of targets not seen. For this reason, the first question addressed was what working memory model and their associated assumption could be extended to the current study’s orientation perception task? Based on goodness-of-fit metrics, we found that the standard mixture model fit the current data set better than the models that assumed a varying distribution for the standard deviation parameter. In other words, we did not find evidence supporting the idea that precision can be described by a variable distribution during orientation perception. This result is consistent with Shen and Ma (2019) who also found little evidence for variable precision during visual perception. Furthermore, we found that the models that included a guess rate parameter performed better than the variable precision models that did not. This indicates that the existence of guessing cannot only be attributed to the low perceptual quality that was drawn, by chance, from a stochastic distribution. It is important to point out that this does not mean there is no variability in performance over the course of an experiment. It has been noted by many authors that performance changes as participants get better at the task or when they get tired towards the end of the experiment. Rather, these results suggest that a model with a fixed standard deviation fits better to orientation perception task performance than one that varies from trial to trial according to some underlying distribution.

### EEG Activity, Model Parameters and Perceptual Behavior

The second goal was to try to answer the question of how brain activity modulates perceptual representation as it is quantified by the guess rate and standard deviation parameters. Specifically, we wanted to see if brain activity prior to the target onset in the 8-14 Hz frequency range (alpha band) modulated whether a target is perceived (*i.e*., guess rate); and, we tested whether the quality of target’s perceptual representation (*i.e*., standard deviation parameter) was related to post-target brain activity in the 4-7 Hz frequency range (theta band). To this end, trials were categorized as accurate or guesses based on model parameter values. The EEG activity on accurate and guess trials were then compared across participants. Then, a complementary approach was used that did a median split of trials based on EEG power. The model was fit to the high power and low power trials and the resulting parameter values were compared across time within different frequency bands. Despite our original hypothesis that alpha activity (8-14 Hz) would be related to guess rate or the standard deviation parameter, the evidence did not support this idea. Instead, the model-based and power-based approaches both found that 2-7 Hz activity after target onset modulated perceptual representation as it was quantified by the guess rate and standard deviation parameters.

#### Accurate vs Guess Trials: ERP Activity

Significant differences in ERP waveforms of the accurate and guess trials started at around 200 ms post-target onset in the anterior locations on the head (Figure 3B) which corresponds to the P2 component (Di Russo et al., 2019; Key et al., 2005; Potts & Tucker, 2001). The greater amplitude of the P2 for accurate guess trails fits with the P2 being related to the salience of stimuli (Potts & Tucker, 2001) or stimulus recognition (Harel et al., 2016).

We also observed a more negative N2 for guess trials, that seems to go against current literature which suggests that the onset of visual consciousness can be marked by a greater negativity in the N2 time range due to an overlapping posterior negative component called the visual awareness negativity (VAN) (Förster et al., 2020; Koivisto & Revonsuo, 2003, 2010). However, the separation of trials based on response performance rather than awareness (according to Koivisto and Revonsuo (2010), the VAN can only be reliably detected by comparing aware and unaware conditions), would explain why we would not expect greater negativity in the N2 time range for accurate ERPs compared to guess ERPs. Furthermore, the distribution of the observed N2 component (mostly right central electrodes) does not match the typical posterior VAN distribution (Förster et al., 2020). In fact, the N2 distribution points towards the presence of the much larger P3 component. The most likely explanation is that the attenuated guess trial ERPs (compared to accurate trials) results in a smaller or later positive voltage from the much larger P3 overlapping the negative deflection at the N2 time period (Förster et al., 2020; Koivisto & Revonsuo, 2010) so that the N2 appears more negative in guess trials when it is actually due to the P3 being smaller. The voltage increase of accurate compared to guess ERPs during the P3 time window follow previous findings on attention and visual perception (Key et al., 2005; Salti et al., 2012). In the current study the P3 is likely related to processes such as stimulus classification and saliency evaluation (Doradziń Knight, 2019; Salti et al., 2012) and that these processes are overwhelmingly more relevant to accurate orientation perception than processes for conscious perception as reflected by the relatively smaller N2 ERP. Furthermore, the difference in ERPs between accurate trials and guess trials during the N2 time period (see middle topographic plot of Figure 3B) indicates that the processes reflected by the N2 likely occur in both types of trials though to differing degrees. Instead, it appears that the main difference between accurate and guess trials is the extent that the target can be classified or evaluated by the brain regardless of consciously perceiving the target.

#### Accurate vs Guess Trials: EEG Activity

We had originally hypothesized activity within the alpha frequency band (8-14 Hz range), especially prior to target onset, mediating task performance. However, the results did not support this hypothesis. It is possible a null effect could be attributed to using both pre- and post-target activity in the pixel-based multiple correction procedure. Relative to post-target activity, pre-stimulus power would be very small since the only thing for participants to do was fixate on a white circle. As a result, differences in pre-target activity might not exceed significance thresholds when those thresholds were based on the background spectra of the combined pre- and post-target activity. However, we tested this possibility by analyzing power in the 8-14 Hz frequency range during just the pre-target time period and found no significant differences between accurate and guess trials (data not shown). Therefore, it is unlikely that a lack of significant difference in pre-target 8-14 Hz range is due to an exceedingly high statistical threshold set by the post-target activity.

Another possibility is that the short intertrial interval (ITI) between participants’ response and fixation onset starting the next trial meant alpha desynchronization carried over to the next trial so that it masked any true effects of spontaneous alpha. While this is a possibility, alpha activity following the end of a trial is also affected by evaluating their response, task load, and motivational state (Compton et al., 2011, 2014, 2017). While the short ITI is problematic for alpha analysis in some regards, it is beneficial in other ways such as discouraging participants from strategizing or reflecting back on the previous trial.

Similarly, the interstimulus interval (ISI) between fixation onset and target onset is possibly too short to let event-related desynchronization (ERD) of alpha to recover, thus masking the effects of spontaneous alpha during the pre-target period. Although potentially problematic, this should not adversely affect results because the ERD should be equal across all conditions since fixation is the same on all trials. Therefore, the activity related to the target perception is equally affected so relative differences can still be detected if they are present.

The most likely reason for not finding a pre-target effect of 8-14 Hz activity is that there was no information for the participant to use before a target was presented, so there would be no attention-related changes or task-based preparation prior to the target. This means that the only activity present during fixation are the spontaneous fluctuations of normal neural activity. Recent work by Samaha et al. (2020) propose that spontaneous alpha activity modulates neural activity in a non-specific manner, thus neither facilitating nor inhibiting perceptual activity. The net effect would be no change to performance which is line with what the model proposed by Samaha and colleagues (2020) would predict.

The absence of significant effects in post-target 8-14 Hz (alpha band) activity suggests that alpha activity is not related to the processes responsible for differences between accurate and guess trials. The fact that post-target 8-14 Hz activity was also unrelated to response errors on a trial-by-trial basis suggests that 8-14 Hz activity is not related to perception of a target’s orientation. This is consistent with Bae and Luck (2018) who found that alpha activity did not encode a target’s orientation when the target’s spatial location was controlled for. An absence of a relationship between 8-14 Hz power and performance measures has been noted in other studies though they were not able to rule out changes in detection bias which can be ruled out in the current study since guess rate was not related to alpha activity (Benwell et al., 2017; Keitel et al., 2018). However, awareness bias might be a possibility assuming orientation perception can proceed in the absence of conscious awareness which seems likely based on previous research (Benwell et al., 2017; Doradzi ska et al., 2020; Koenig & Ro, 2019). In sum, it is likely that 8-14 Hz (alpha band) activity is more related to a global process that is not dependent on conscious awareness and would not change across trials such as feature-independent stimulus processing.

The most significant differences between accurate and guess trials were in the 2-4 Hz frequency ranges, particularly in the parietal and parietooccipital electrodes during the latter half of the post-stimulus epoch (Karakaş, 2020). A significant increase of delta and low theta power in accurate compared to guess trials could be seen at around 300 ms though their differences in mean phase, especially for theta, started earlier (200 ms post-target). Although there was a trend in our data for increased theta power in accurate trials compared to guess trials, this difference was not reliable.

The significant differences in theta and delta seem to correspond to changes in the N2 and P3 ERP components observed in the time domain. Many studies have shown that delta and theta are the primary contributors to the formation of the N2 and P3 (Harmony, 2013; Harper et al., 2014; Karakaş et al., 2000b). It has been proposed that activity in the 2-3 Hz range contributes continuous positivity throughout the ERP response while theta activity corresponds to the polarity change as the negative deflection during the N2 shifts to the positive deflection during the P3 (Harper et al., 2014). This fits with the observed EEG activity of accurate and guess trials (Figure 3C and 3D). Furthermore, the interplay between theta and delta activity would explain the indistinct N2 but large P3 waveforms (Karakaş et al., 2000b).

The increase in baseline normalized 4-7 Hz power during accurate trials relative to guess trials has the earliest onset in left frontal and right parietal electrodes. Prolonged theta in right posterior electrodes is accompanied by significant differences in mean (low) theta phase between accurate and guess trials. Theta has been interpreted as being correlated with selective attentional processing (Başar et al., 2001; Karaka , 2020; Karakaş et al., 2000b) and an increase in theta synchronization and power has been associated with successful memory encoding (Klimesch, 1999) and right hemispheric theta is greater than left when encoding visuospatial information (Sauseng et al., 2004). Although the task in this study was not a memory task, encoding information is still a viable way of looking at how the brain transforms the visual sensory information into perceptual representations. In fact, theta has been shown to be sensitive to target and non-target stimuli regardless of memory load (Palomäki et al., 2012). Therefore, it is probable that theta activity might be related to the “encoding” of the target’s orientation or just the detectability of target itself. Both functions could occur in accurate and guess trials, though more often or to a greater degree in accurate trials compared to guess trials. This would explain why 4-7 Hz activity showed a for trend increased power in accurate trials compared to guess trials though this difference was usually not significant.

In contrast to the 4-7 Hz activity, activity within 2-3 Hz showed a more spatially diffuse increase in power in accurate vs guess trials with maxima over the lateral parietal areas. Delta phase differed significantly between accurate and guess trials over the occipital and lateral parietal areas with the most robust phase differences at around 350 ms post-target. Several experiments that showed increases in delta activity during the performance of different mental tasks and conflict-monitoring paradigms have led investigators to associate delta with modulation of networks that should be inactive to accomplish the task (Harmony, 2013; Rawls et al., 2020). Investigators have also noted a close association between delta activity and the P3 ERP waveform (Harper et al., 2014; Rawls et al., 2020; Schürmann et al., 1995, 2001). If the P3 reflects post-perceptual processes, then it is likely 2-3 Hz activity is also involved. Whether that role is as an inhibitory mechanism or not remains unknown. Overall, when considering the time and frequency at which reliable differences were observed between accurate and guess trials, it seems likely that this post-target activity reflects differences in the level of perceptual encoding of the target orientation and the subsequent precision of responses on the task.

#### Single-Trial EEG Activity and Response Errors

We were surprised to find no significant correlation between response error and oscillatory power in any of the measured frequencies, but this could also be attributed to the lack of sensitivity of the analysis method. Since failed to find support for our hypothesis regarding alpha oscillations, we chose to take a conservative approach when analyzing the rest of the time-frequency space. Using a less conservative method might help answer the question but that would increase the risk of finding false positives which could be more harmful than accepting false negatives.

Interestingly, we did find a relationship between response error and phase values in the 2- 7 Hz frequencies similar to the significant differences in mean phase of accurate trials compared to guess trials. In fact, phase modulation by response errors is even more pronounced than the differences in phase between accurate and guess trials. This strongly suggests 2-7 Hz phase activity plays an important role in the amount of participants’ response error on a trial-by-trial bases. Delta (2-3 Hz) and theta (4-7 Hz) frequencies are usually associated with working memory and cognitive control, but these results imply that they have an important role in visual perceptual processes as well.

#### Relationship Between Log Power and Standard Mixed Model Parameters

There were significant differences in the guess rate and standard deviation parameter values from trials with high compared to low log power at 2-3 Hz starting at around 255 ms post-target onset. The trend was for trials with low power to have higher guess rate and standard deviation parameter values than trials with high power. The same trend was found for the guess rates on trials with high vs low 4-7 Hz log power though those effects started a little earlier (around 200 ms post-target onset) than those at 2-3 Hz frequency range. These results are in line with the accurate vs guess analysis performed on baseline normalized power data. Specifically, trials categorized as “accurate” had more power and a different preferred phase in the 2-7 Hz frequency range than trials considered “guesses.” There were no significant effects prior to the target onset which fits with the baseline sensory excitability model proposed by Samaha and colleagues (2020).

Interestingly, Fp1 electrode showed significant difference in guess rate between high and low power trials prior to the onset of the response screen. Accumulation of evidence has led investigators to associate frontal delta (2-3 Hz) activity with top-down control and response inhibition (De Vries et al., 2018; Harmony, 2013; Helfrich et al., 2017; Rawls et al., 2020).

Considering that these late significant effects in the frontal area are almost entirely after those in the parietal and occipital electrodes and are only for guess rate, it is likely top-down control has to do with maintaining the perceptual representation or inhibiting distracting information during the delay period. In this way, an increase in 2-3 Hz brain activity would help maintain a target’s “perceived” state rather than the target becoming part of the “unseen” distribution that the guess rate represents.

Considering the timing and frequency of these significant effects, it seems likely that they reflect the same processes measured by the ERP components in the accurate vs guess analysis. However, as others have found (and can be seen in Figure 8 and Tables 2 and 3), an ERP component’s amplitude is usually determined by a combination of different underlying brain potentials which often represent separable functional processes (Harper et al., 2014; Karakaş et al., 2000a; Woodman, 2010). In the current study, it is likely that the activity at 2-3 Hz and 4-7 Hz interact dynamically to contribute to the measured ERP components morphology.

### Conclusion and Future Directions

Overall, we have shown that the standard mixture model can be extended to a visual perception task and that doing so provides important insight into how brain activity shapes visual perception. Our results point towards a perceptual representation that has a fixed precision meaning that while response errors vary with the level of neural activity after stimulus onset, their variability is according to the same distribution across trials. Whether this is true across different levels of target visibility is not known but would be an interesting question for future research.

It is important to note that our results are not suggesting pre-target EEG activity has no effect on visual perception. There is a lot of evidence to contrary. It has been proposed that pre-target alpha activity alters participants’ confidence rather than performance on visual perception tasks (Samaha et al., 2017, 2020), a distinction the current study was not designed to test. It is possible the analysis methods were not sensitive enough to pick up changes in pre-target activity though there were no significant effects in a follow-up analysis analyzing just the pre-target time period making a lack of statistical power unlikely. The most likely explanation for an absence of pre-target effects is that participants did not have information about an upcoming trial so there was no reason to prepare prior to the target. A follow-up study that directly manipulates attention or detectability of the target and measures participants’ confidence along with the objective measures of response error, might be able to better test how differences in brain activity prior to target onset affect visual perception and subsequent task performance.

## Acknowledgements

This research was supported by start-up funds from the Faculty of Science and an NSERC Discovery grant (#RES0024267) awarded to Kyle Elliot Mathewson. The authors would like you thank Erron Jacob Meneses, Jenny Le, and David Lam for assistance in data collection, and Daniel Robles and Jonathan Kuziek for analysis feedback.

## Author Contributions

KEM and SSS conceived the experiment. SSS collected and analyzed the data. KEM facilitated the results interpretation. SSS wrote the first draft of the manuscript. SSS and KEM edited the manuscript.

## Data Accessibility

The data that support the findings of this study are openly available through an Open Science Framework repository at https://doi.org/10.17605/OSF.IO/CW7UX. Scripts to reproduce all reported analyses and figures are available through a Github repository at https://github.com/APPLabUofA/OrientTask_paper.

