## Supplementary figures and images for "To See, Not to See, or to See Poorly: Perceptual Quality and Guess Rate as a Function of Electroencephalography (EEG) Brain Activity in an Orientation Perception Task"

### Figure S1

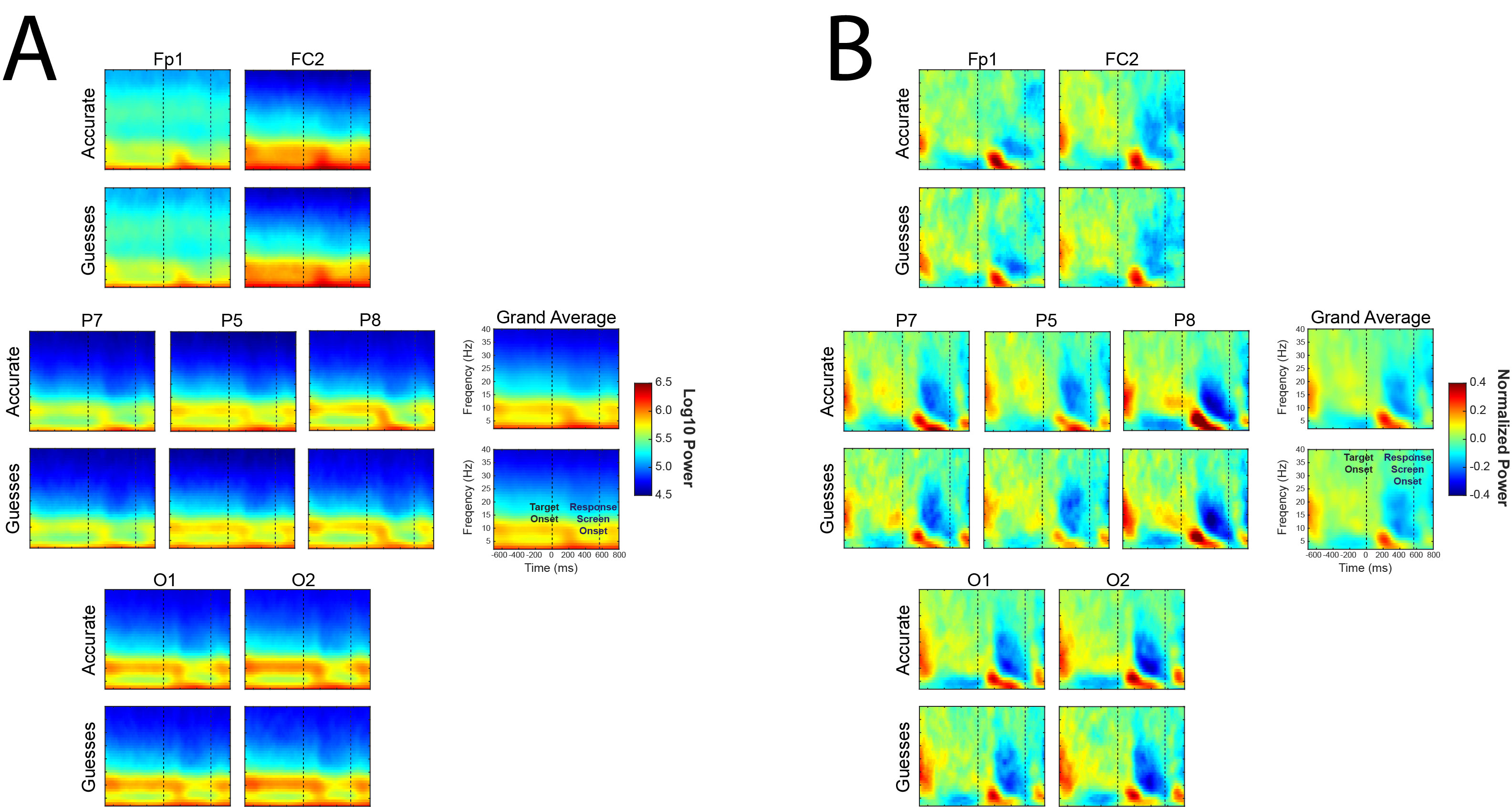
