## Supporting Information for "To See, Not to See, or to See Poorly: Perceptual Quality and Guess Rate as a Function of Electroencephalography (EEG) Brain Activity in an Orientation Perception Task"

### Methods

Stepwise multiple regression analyses were performed for the standard mixture model parameters from “high” and “low” trials separately. For the guess rate parameter, predictor variables were the guess rates on 2-3 Hz, 4-7 Hz, and 8-40 Hz log power trials averaged across the time windows used for each ERP component that showed significant effects in the previous analyses: 200-255 ms (P2), 255-360 ms (N2), and 360-500 ms (P3). The predicted variables were the guess rates on trials with high or low amplitudes in their P2, N2, and P3 ERP components. For the standard deviation parameter, the predictor variables were the same in the guess rate analyses except only the 360-500 ms (P3) time window was used. Similarly, the only predicted variables were the standard deviation parameters from trials split by their P3 ERP component. A Bonferroni type adjustment was made for inflated Type 1 error, α for each electrode site was assigned the value of 0.0028 for each *b** among a set of *b** such that α for each set did not exceed the critical value of 0.05 (Karakaş et al., 2000) The cumulative proportion of explained variances adjusted for number of predictors (referred to as the adjusted coefficient of determination; $R_{adj}^{2}$) and the standardized regression coefficients (*b**) for the regression equations at each step of the multivariate analysis for guess rates and standard deviation parameters are summarized in Table 2 and Table 3, respectively.

It should be noted that multicollinearity, that is, a high correlation between two or most predictor variables, is present in the stepwise multiple regression analyses described above. This is the most likely cause of the negative standardized regression coefficients seen in Tables 2 and 3. While this is considered an issue that could interfere with the multiple regression analyses, we are using these results to get a rough estimate of the relative contributions from each predictor variable. The presence of multicollinearity usually means that it is harder to reject the null hypothesis, thus one might miss the importance of the predictors (Faraway, 2016). With regards to the current dataset, this means that some predictors might have greater relative importance than indicated by their *b**.

### Results

As Table S2 shows, the guess rate parameter from 2-3 Hz, 4-7 Hz and higher frequencies amply account for the guess rate values on trials with high and low ERP amplitudes. The proportion of explained variances were between 0.935-0.990. Table 3 shows a similar result for the standard deviation (σ) parameter except the proportion of explained variances were smaller, being between 0.700-0.928.

The guess rate from the P2 (200-255 ms) ERP trials (both high and low) had a clear difference in its major contributor between the frontal electrodes (Fp1 and FC2) and the parietal and occipital electrodes. The guess rates from 2-3 Hz and 2-7 Hz trials accounted for most of the variance (0.954-0.976) in the frontal electrodes and the addition of higher frequencies was redundant or detrimental (0.952-0.975). In comparison, the addition of higher frequencies in the parietal and occipital electrodes regression models added some small benefit to the proportion of explained variances (increased 0.009-0.036).

Interestingly, the guess rate from the N2 (255-360 ms) ERP trials (both high and low) had most of its variance accounted for by the guess rates from 2-3 Hz trials (0.955-0.982). According to the standardized regression coefficients, whose values can be thought of as indicators of relative importance in the regression model, the 2-3 Hz guess rates were the “best” predictors in all cases except for Fp1 which indicates 4-7 Hz as the best predictor, and O2 which has the 8-40 Hz variable as the best predictor.

Unlike the other ERPs, guess rate from the P3 (360-500 ms) ERP trials had a difference in the proportion of explained variances between the high and low trials. That is, at all electrodes, the trials categorized as “low” had more explained variance by the set of predictors than trials categorized as “high” (0.964-0.980 and 0.935-0.976, respectively). In all cases, the majority of the proportion of explained variances came from the 2-3 Hz guess rates though 4-7 Hz was often a relatively important predictor. The exceptions were O1 and O2 which consistently had 8-40 Hz as the most important predictor variable.

Like the results from the guess rate analyses of the P3 (360-500 ms) ERP trials, the standard deviation (σ) parameter indicated that the proportion of explained variances differed between trials categorized as “low” compared to “high.” However, Fp1 and FC2 had more explained variance for the low category of trials than the high (see Table S3), whereas the parietal and occipital electrodes were the reverse (low: 0.700-0.898; high: 0.867-0.928). Interestingly, the most important predictor variables tended to differ between the high and low categories. Most of the time, 4-7 Hz was ranked the most important predictor in the high category, whereas 2-3 Hz was ranked the most important predictor in the low category. The notable exception was the low category of the O2 electrode which had the proportion of explained variance increase from 0.570 to 0.709 with the addition of the 4-7 Hz predictor.

### Tables

**Table S1.**

*Fitted parameter values for the standard mixture model with and without a bias parameter, and the two variable precision models with and without a guess rate parameter.*

|  | Standard Mixture Model | | |
| --- | --- | --- | --- |
|  | Guess Rate (g) | SD (σ) | Bias (µ) |
| Model | 0.19 ± 0.04 | 11.15 ± 0.51 | - |
| Model + Bias | 0.19 ± 0.04 | 11.04 ± 0.51 | 0.14 ± 0.29 |
|  | Variable Precision Model (Gaussian over SD) | | |
|  | Guess Rate (g) | meanSD (σ_mn_) | stdSD (σ_std_) |
| Model | 0.18 ± 0.03 | 11.26 ± 0.54 | 2.82 ± 0.31 |
| Model − Guess Rate | - | 10.23 ± 5.72 | 33.48 ± 5.26 |
|  | Variable Precision Model (Gamma over Precision) | | |
|  | Guess Rate (g) | modePrecision (J_mod_) / Shape | stdPrecision (J_std_) / Scale |
| Model | 0.17 ± 0.03 | 0.012 ± 0.001 / 13.62 ± 5.42 | 0.009 ± 0.002 / 0.005 ± 0.001 |
| Model − Guess Rate | - | 0.011 ± 0.002 / 1.59 ± 0.12 | 0.061 ± 0.017 / 0.056 ± 0.018 |

Note. All values are *Mean* ± *SEM*. Scale and shape are the traditional parameters of the Gamma distribution.

**Table S2**

*Stepwise regression results for guess rate (g) parameter.*

| Trial Category | Predicted | P2 (200-255 ms) | |  | N2 (255-360 ms) | |  | P3 (360-500 ms) | |
| --- | --- | --- | --- | --- | --- | --- | --- | --- | --- |
|  | Model | *b** | $R_{adj}^{2}$ |  | *b** | $R_{adj}^{2}$ |  | *b** | $R_{adj}^{2}$ |
| Fp1 | | | | | | | | | |
| High | Step 1 |  | 0.953 |  |  | 0.961 |  |  | 0.958 |
|  | 2-3 Hz | 0.98 |  |  | 0.98 |  |  | 0.98 |  |
|  | Step 2 |  | 0.956 |  |  | 0.965 |  |  | 0.956 |
|  | 2-3 Hz | 0.34 |  |  | 0.29 |  |  | 0.90 |  |
|  | 4-7 Hz | 0.64 |  |  | 0.70 |  |  | 0.08 |  |
|  | Step 3 |  | 0.953 |  |  | 0.963 |  |  | 0.955 |
|  | 2-3 Hz | 0.35 |  |  | 0.35 |  |  | 1.06 |  |
|  | 4-7 Hz | 0.58 |  |  | 0.81 |  |  | 0.24 |  |
|  | 8-40 Hz | 0.06 |  |  | -0.17 |  |  | -0.32 |  |
| Low | Step 1 |  | 0.963 |  |  | 0.970 |  |  | 0.966 |
|  | 2-3 Hz | 0.98 |  |  | 0.99 |  |  | 0.98 |  |
|  | Step 2 |  | 0.966 |  |  | 0.974 |  |  | 0.965 |
|  | 2-3 Hz | 0.31 |  |  | 0.26 |  |  | 0.74 |  |
|  | 4-7 Hz | 0.67 |  |  | 0.73 |  |  | 0.25 |  |
|  | Step 3 |  | 0.965 |  |  | 0.973 |  |  | 0.964 |
|  | 2-3 Hz | 0.30 |  |  | 0.29 |  |  | 0.94 |  |
|  | 4-7 Hz | 0.58 |  |  | 0.75 |  |  | 0.38 |  |
|  | 8-40 Hz | 0.11 |  |  | -0.05 |  |  | -0.34 |  |
| FC2 | | | | | | | | | |
| High | Step 1 |  | 0.953 |  |  | 0.977 |  |  | 0.935 |
|  | 2-3 Hz | 0.98 |  |  | 0.99 |  |  | 0.97 |  |
|  | Step 2 |  | 0.954 |  |  | 0.977 |  |  | 0.935 |
|  | 2-3 Hz | 0.67 |  |  | 1.16 |  |  | 0.67 |  |
|  | 4-7 Hz | 0.31 |  |  | -0.18 |  |  | 0.30 |  |
|  | Step 3 |  | 0.952 |  |  | 0.980 |  |  | 0.935 |
|  | 2-3 Hz | 0.70 |  |  | 0.99 |  |  | 0.50 |  |
|  | 4-7 Hz | 0.41 |  |  | -0.57 |  |  | 0.09 |  |
|  | 8-40 Hz | -0.13 |  |  | 0.56 |  |  | 0.38 |  |
| Low | Step 1 |  | 0.975 |  |  | 0.982 |  |  | 0.965 |
|  | 2-3 Hz | 0.99 |  |  | 0.99 |  |  | 0.98 |  |
|  | Step 2 |  | 0.976 |  |  | 0.983 |  |  | 0.967 |
|  | 2-3 Hz | 0.70 |  |  | 1.23 |  |  | 0.63 |  |
|  | 4-7 Hz | 0.29 |  |  | -0.24 |  |  | 0.36 |  |
|  | Step 3 |  | 0.975 |  |  | 0.984 |  |  | 0.970 |
|  | 2-3 Hz | 0.71 |  |  | 1.07 |  |  | 0.36 |  |
|  | 4-7 Hz | 0.33 |  |  | -0.49 |  |  | 0.05 |  |
|  | 8-40 Hz | -0.05 |  |  | 0.41 |  |  | 0.58 |  |
| P5 | | | | | | | | | |
| High | Step 1 |  | 0.954 |  |  | 0.972 |  |  | 0.963 |
|  | 2-3 Hz | 0.98 |  |  | 0.99 |  |  | 0.98 |  |
|  | Step 2 |  | 0.965 |  |  | 0.971 |  |  | 0.964 |
|  | 2-3 Hz | 0.31 |  |  | 1.09 |  |  | 0.70 |  |
|  | 4-7 Hz | 0.68 |  |  | -0.10 |  |  | 0.29 |  |
|  | Step 3 |  | 0.990 |  |  | 0.972 |  |  | 0.962 |
|  | 2-3 Hz | 0.23 |  |  | 1.04 |  |  | 0.70 |  |
|  | 4-7 Hz | -0.44 |  |  | -0.42 |  |  | 0.28 |  |
|  | 8-40 Hz | 1.21 |  |  | 0.37 |  |  | 0.01 |  |
| Low | Step 1 |  | 0.950 |  |  | 0.980 |  |  | 0.981 |
|  | 2-3 Hz | 0.98 |  |  | 0.99 |  |  | 0.99 |  |
|  | Step 2 |  | 0.960 |  |  | 0.981 |  |  | 0.981 |
|  | 2-3 Hz | 0.29 |  |  | 1.23 |  |  | 0.79 |  |
|  | 4-7 Hz | 0.70 |  |  | -0.24 |  |  | 0.20 |  |
|  | Step 3 |  | 0.986 |  |  | 0.980 |  |  | 0.980 |
|  | 2-3 Hz | 0.20 |  |  | 1.22 |  |  | 0.79 |  |
|  | 4-7 Hz | -0.41 |  |  | -0.30 |  |  | 0.19 |  |
|  | 8-40 Hz | 1.20 |  |  | 0.07 |  |  | 0.01 |  |
| P7 | | | | | | | | | |
| High | Step 1 |  | 0.943 |  |  | 0.968 |  |  | 0.955 |
|  | 2-3 Hz | 0.97 |  |  | 0.98 |  |  | 0.98 |  |
|  | Step 2 |  | 0.956 |  |  | 0.967 |  |  | 0.958 |
|  | 2-3 Hz | 0.05 |  |  | 0.85 |  |  | 0.44 |  |
|  | 4-7 Hz | 0.93 |  |  | 0.13 |  |  | 0.55 |  |
|  | Step 3 |  | 0.988 |  |  | 0.966 |  |  | 0.957 |
|  | 2-3 Hz | -0.13 |  |  | 0.84 |  |  | 0.48 |  |
|  | 4-7 Hz | -0.11 |  |  | 0.03 |  |  | 0.71 |  |
|  | 8-40 Hz | 1.23 |  |  | 0.12 |  |  | -0.21 |  |
| Low | Step 1 |  | 0.942 |  |  | 0.971 |  |  | 0.961 |
|  | 2-3 Hz | 0.97 |  |  | 0.99 |  |  | 0.98 |  |
|  | Step 2 |  | 0.950 |  |  | 0.970 |  |  | 0.967 |
|  | 2-3 Hz | 0.25 |  |  | 1.02 |  |  | 0.09 |  |
|  | 4-7 Hz | 0.73 |  |  | -0.04 |  |  | 0.89 |  |
|  | Step 3 |  | 0.987 |  |  | 0.969 |  |  | 0.967 |
|  | 2-3 Hz | -0.02 |  |  | 0.96 |  |  | 0.23 |  |
|  | 4-7 Hz | -0.27 |  |  | -0.21 |  |  | 1.11 |  |
|  | 8-40 Hz | 1.29 |  |  | 0.25 |  |  | -0.36 |  |
| P8 | | | | | | | | | |
| High | Step 1 |  | 0.952 |  |  | 0.955 |  |  | 0.964 |
|  | 2-3 Hz | 0.98 |  |  | 0.98 |  |  | 0.98 |  |
|  | Step 2 |  | 0.950 |  |  | 0.954 |  |  | 0.975 |
|  | 2-3 Hz | 1.12 |  |  | 0.72 |  |  | 0.15 |  |
|  | 4-7 Hz | -0.15 |  |  | 0.26 |  |  | 0.84 |  |
|  | Step 3 |  | 0.973 |  |  | 0.960 |  |  | 0.976 |
|  | 2-3 Hz | 0.39 |  |  | 0.52 |  |  | -0.06 |  |
|  | 4-7 Hz | -0.77 |  |  | -0.44 |  |  | 0.62 |  |
|  | 8-40 Hz | 1.36 |  |  | 0.90 |  |  | 0.43 |  |
| Low | Step 1 |  | 0.943 |  |  | 0.967 |  |  | 0.982 |
|  | 2-3 Hz | 0.97 |  |  | 0.98 |  |  | 0.99 |  |
|  | Step 2 |  | 0.941 |  |  | 0.966 |  |  | 0.983 |
|  | 2-3 Hz | 1.13 |  |  | 0.85 |  |  | 0.64 |  |
|  | 4-7 Hz | -0.16 |  |  | 0.14 |  |  | 0.35 |  |
|  | Step 3 |  | 0.964 |  |  | 0.969 |  |  | 0.984 |
|  | 2-3 Hz | 0.27 |  |  | 0.62 |  |  | 0.32 |  |
|  | 4-7 Hz | -0.70 |  |  | -0.45 |  |  | 0.23 |  |
|  | 8-40 Hz | 1.41 |  |  | 0.82 |  |  | 0.44 |  |
| O1 | | | | | | | | | |
| High | Step 1 |  | 0.959 |  |  | 0.958 |  |  | 0.950 |
|  | 2-3 Hz | 0.98 |  |  | 0.98 |  |  | 0.98 |  |
|  | Step 2 |  | 0.957 |  |  | 0.956 |  |  | 0.956 |
|  | 2-3 Hz | 0.76 |  |  | 1.05 |  |  | 0.47 |  |
|  | 4-7 Hz | 0.22 |  |  | -0.07 |  |  | 0.51 |  |
|  | Step 3 |  | 0.967 |  |  | 0.958 |  |  | 0.958 |
|  | 2-3 Hz | 0.29 |  |  | 0.80 |  |  | 0.29 |  |
|  | 4-7 Hz | -0.13 |  |  | -0.31 |  |  | 0.21 |  |
|  | 8-40 Hz | 0.83 |  |  | 0.49 |  |  | 0.49 |  |
| Low | Step 1 |  | 0.957 |  |  | 0.967 |  |  | 0.972 |
|  | 2-3 Hz | 0.98 |  |  | 0.98 |  |  | 0.99 |  |
|  | Step 2 |  | 0.956 |  |  | 0.966 |  |  | 0.975 |
|  | 2-3 Hz | 0.71 |  |  | 1.10 |  |  | 0.53 |  |
|  | 4-7 Hz | 0.27 |  |  | -0.12 |  |  | 0.46 |  |
|  | Step 3 |  | 0.967 |  |  | 0.971 |  |  | 0.982 |
|  | 2-3 Hz | 0.26 |  |  | 0.90 |  |  | 0.28 |  |
|  | 4-7 Hz | -0.09 |  |  | -0.64 |  |  | -0.18 |  |
|  | 8-40 Hz | 0.82 |  |  | 0.72 |  |  | 0.90 |  |
| O2 | | | | | | | | | |
| High | Step 1 |  | 0.932 |  |  | 0.956 |  |  | 0.954 |
|  | 2-3 Hz | 0.97 |  |  | 0.98 |  |  | 0.98 |  |
|  | Step 2 |  | 0.945 |  |  | 0.954 |  |  | 0.960 |
|  | 2-3 Hz | 0.01 |  |  | 0.98 |  |  | 0.36 |  |
|  | 4-7 Hz | 0.97 |  |  | 0.00 |  |  | 0.62 |  |
|  | Step 3 |  | 0.964 |  |  | 0.969 |  |  | 0.963 |
|  | 2-3 Hz | -0.23 |  |  | 0.41 |  |  | 0.14 |  |
|  | 4-7 Hz | -0.06 |  |  | -0.54 |  |  | 0.27 |  |
|  | 8-40 Hz | 1.28 |  |  | 1.12 |  |  | 0.58 |  |
| Low | Step 1 |  | 0.910 |  |  | 0.960 |  |  | 0.972 |
|  | 2-3 Hz | 0.96 |  |  | 0.98 |  |  | 0.99 |  |
|  | Step 2 |  | 0.938 |  |  | 0.960 |  |  | 0.975 |
|  | 2-3 Hz | -0.17 |  |  | 0.74 |  |  | 0.52 |  |
|  | 4-7 Hz | 1.14 |  |  | 0.24 |  |  | 0.47 |  |
|  | Step 3 |  | 0.959 |  |  | 0.972 |  |  | 0.979 |
|  | 2-3 Hz | -0.37 |  |  | 0.25 |  |  | 0.25 |  |
|  | 4-7 Hz | 0.10 |  |  | -0.44 |  |  | 0.10 |  |
|  | 8-40 Hz | 1.25 |  |  | 1.18 |  |  | 0.64 |  |
| *Note.* Predictors in each model are the guess rate parameters from fitting the standard mixture model to trials categorized as high or low log power in the specified frequency band and time windows. Predicted values are the guess rate parameters from fitting the standard mixture model to trials categorized as high or low ERP amplitudes in the specified time windows. *b** = estimated values of standardized regression coefficients; $R_{adj}^{2}$ = adjusted coefficient of determination (*i.e.,* coefficient of determination adjusted for number of predictors). | | | | | | | | | |

**Table S3**

*Stepwise regression results for the standard deviation (σ) parameter.*

| Trial Category | Predicted | P3 (360-500 ms) | |
| --- | --- | --- | --- |
|  | Model | *b** | $R_{adj}^{2}$ |
| Fp1 | | | |
| High | Step 1 |  | 0.865 |
|  | 2-3 Hz | 0.93 |  |
|  | Step 2 |  | 0.904 |
|  | 2-3 Hz | 0.34 |  |
|  | 4-7 Hz | 0.63 |  |
|  | Step 3 |  | 0.899 |
|  | 2-3 Hz | 0.29 |  |
|  | 4-7 Hz | 0.56 |  |
|  | 8-40 Hz | 0.11 |  |
| Low | Step 1 |  | 0.884 |
|  | 2-3 Hz | 0.94 |  |
|  | Step 2 |  | 0.916 |
|  | 2-3 Hz | 0.64 |  |
|  | 4-7 Hz | 0.35 |  |
|  | Step 3 |  | 0.912 |
|  | 2-3 Hz | 0.65 |  |
|  | 4-7 Hz | 0.37 |  |
|  | 8-40 Hz | -0.02 |  |
| FC2 | | | |
| High | Step 1 |  | 0.879 |
|  | 2-3 Hz | 0.94 |  |
|  | Step 2 |  | 0.880 |
|  | 2-3 Hz | 0.75 |  |
|  | 4-7 Hz | 0.21 |  |
|  | Step 3 |  | 0.906 |
|  | 2-3 Hz | 0.24 |  |
|  | 4-7 Hz | 0.09 |  |
|  | 8-40 Hz | 0.65 |  |
| Low | Step 1 |  | 0.893 |
|  | 2-3 Hz | 0.95 |  |
|  | Step 2 |  | 0.913 |
|  | 2-3 Hz | 0.53 |  |
|  | 4-7 Hz | 0.44 |  |
|  | Step 3 |  | 0.915 |
|  | 2-3 Hz | 0.34 |  |
|  | 4-7 Hz | 0.35 |  |
|  | 8-40 Hz | 0.29 |  |
| P5 | | | |
| High | Step 1 |  | 0.921 |
|  | 2-3 Hz | 0.96 |  |
|  | Step 2 |  | 0.931 |
|  | 2-3 Hz | 0.42 |  |
|  | 4-7 Hz | 0.55 |  |
|  | Step 3 |  | 0.928 |
|  | 2-3 Hz | 0.51 |  |
|  | 4-7 Hz | 0.60 |  |
|  | 8-40 Hz | -0.13 |  |
| Low | Step 1 |  | 0.787 |
|  | 2-3 Hz | 0.89 |  |
|  | Step 2 |  | 0.791 |
|  | 2-3 Hz | 1.33 |  |
|  | 4-7 Hz | -0.46 |  |
|  | Step 3 |  | 0.788 |
|  | 2-3 Hz | 1.79 |  |
|  | 4-7 Hz | -0.42 |  |
|  | 8-40 Hz | -0.49 |  |
| P7 | | | |
| High | Step 1 |  | 0.849 |
|  | 2-3 Hz | 0.93 |  |
|  | Step 2 |  | 0.870 |
|  | 2-3 Hz | 0.02 |  |
|  | 4-7 Hz | 0.92 |  |
|  | Step 3 |  | 0.867 |
|  | 2-3 Hz | 0.24 |  |
|  | 4-7 Hz | 1.09 |  |
|  | 8-40 Hz | -0.39 |  |
| Low | Step 1 |  | 0.721 |
|  | 2-3 Hz | 0.86 |  |
|  | Step 2 |  | 0.722 |
|  | 2-3 Hz | 1.45 |  |
|  | 4-7 Hz | -0.61 |  |
|  | Step 3 |  | 0.738 |
|  | 2-3 Hz | 1.69 |  |
|  | 4-7 Hz | -1.43 |  |
|  | 8-40 Hz | 0.62 |  |
| P8 | | | |
| High | Step 1 |  | 0.911 |
|  | 2-3 Hz | 0.96 |  |
|  | Step 2 |  | 0.913 |
|  | 2-3 Hz | 0.68 |  |
|  | 4-7 Hz | 0.29 |  |
|  | Step 3 |  | 0.924 |
|  | 2-3 Hz | 0.45 |  |
|  | 4-7 Hz | 0.10 |  |
|  | 8-40 Hz | 0.43 |  |
| Low | Step 1 |  | 0.875 |
|  | 2-3 Hz | 0.94 |  |
|  | Step 2 |  | 0.890 |
|  | 2-3 Hz | 0.66 |  |
|  | 4-7 Hz | 0.31 |  |
|  | Step 3 |  | 0.898 |
|  | 2-3 Hz | 0.71 |  |
|  | 4-7 Hz | -0.12 |  |
|  | 8-40 Hz | 0.40 |  |
| O1 | | | |
| High | Step 1 |  | 0.903 |
|  | 2-3 Hz | 0.95 |  |
|  | Step 2 |  | 0.919 |
|  | 2-3 Hz | 0.40 |  |
|  | 4-7 Hz | 0.57 |  |
|  | Step 3 |  | 0.926 |
|  | 2-3 Hz | 0.00 |  |
|  | 4-7 Hz | 0.49 |  |
|  | 8-40 Hz | 0.49 |  |
| Low | Step 1 |  | 0.859 |
|  | 2-3 Hz | 0.93 |  |
|  | Step 2 |  | 0.855 |
|  | 2-3 Hz | 0.81 |  |
|  | 4-7 Hz | 0.12 |  |
|  | Step 3 |  | 0.880 |
|  | 2-3 Hz | 1.54 |  |
|  | 4-7 Hz | 0.01 |  |
|  | 8-40 Hz | -0.64 |  |
| O2 | | | |
| High | Step 1 |  | 0.850 |
|  | 2-3 Hz | 0.93 |  |
|  | Step 2 |  | 0.881 |
|  | 2-3 Hz | 0.43 |  |
|  | 4-7 Hz | 0.52 |  |
|  | Step 3 |  | 0.875 |
|  | 2-3 Hz | 0.37 |  |
|  | 4-7 Hz | 0.51 |  |
|  | 8-40 Hz | 0.07 |  |
| Low | Step 1 |  | 0.570 |
|  | 2-3 Hz | 0.77 |  |
|  | Step 2 |  | 0.709 |
|  | 2-3 Hz | -0.20 |  |
|  | 4-7 Hz | 1.04 |  |
|  | Step 3 |  | 0.700 |
|  | 2-3 Hz | 0.10 |  |
|  | 4-7 Hz | 1.14 |  |
|  | 8-40 Hz | -0.40 |  |
| *Note.* Predictors in each model are the guess rate parameters from fitting the standard mixture model to trials categorized as high or low log power in the specified frequency band and time windows. Predicted values are the standard deviation parameters from fitting the standard mixture model to trials categorized as high or low ERP amplitudes in the specified time windows. *b** = estimated values of standardized regression coefficients; $R_{adj}^{2}$ = adjusted coefficient of determination (*i.e.,* coefficient of determination adjusted for number of predictors). | | | |

**Figure Captions**

**Figure S1.**

**A)** Time-frequency plots showing the mean raw log power of trials categorized as accurate and guesses at selected electrodes and the average of all electrodes (Grand Average). **B)** Time-frequency plots showing the mean power normalized by baseline power of trials categorized as accurate and guesses at selected electrodes and the average of all electrodes (Grand Average).
